# Characterization of Constitutive macro-ER-phagy

**DOI:** 10.1101/2020.10.21.349167

**Authors:** Zhanna Lipatova, Valeriya Gyurkovska, Sarah F. Zhao, Nava Segev

**Affiliations:** Department of Biochemistry and Molecular Genetics, University of Illinois at Chicago

**Keywords:** macro-autophagy, ER-phagy, reticulophagy, Atgs, Ypt1, Ypt/Rab, Atg39, Atg40, ERAD, ERAD-C, Doa10, ERAD-M, Hrd1

## Abstract

Thirty percent of all cellular proteins are inserted into the endoplasmic reticulum (ER), which spans throughout the cytoplasm. Two well-established stress-induced pathways ensure quality control (QC) at the ER: ER-phagy and ER-associated degradation (ERAD), which shuttle cargo for degradation to the lysosome and proteasome, respectively. In contrast, not much is known about constitutive ER-phagy. We have previously reported that excess of integral-membrane proteins is delivered from the ER to the lysosome via autophagy during normal growth of yeast cells. Here, we characterize this pathway as constitutive ER-phagy. Constitutive and stress-induced ER-phagy share the basic macro-autophagy machinery including the conserved Atgs and Ypt1 GTPase. However, induction of stress-induced autophagy is not needed for constitutive ER-phagy to occur. Moreover, the selective receptors needed for starvation-induced ER-phagy, Atg39 and Atg40, are not required for constitutive ER-phagy and neither these receptors nor their cargos are delivered through it to the vacuole. As for ERAD, while constitutive ER-phagy recognizes cargo different from that recognized by ERAD, these two ER-QC pathways can partially substitute for each other. Because accumulation of membrane proteins is associated with disease, and constitutive ER-phagy players are conserved from yeast to mammalian cells, this process could be critical for human health.

**Author Summary:** Accumulation of excess proteins can lead to their aggregation, which in turn can cause multiple disorders, notably neurodegenerative disease. Nutritional and endoplasmic-reticulum (ER) stress stimulate autophagy and ER-associated degradation (ERAD) to clear excess and misfolded proteins, respectively. However, not much is known about clearance of excess proteins during normal growth. We have previously shown that excess integral-membrane proteins are cleared from the ER by macro-autophagy during normal growth of yeast cells. Here we characterize this pathway as constitutive ER-phagy. While this pathway shares machinery of core Atgs and autophagosomes with nutritional stress-induced ER-phagy, it differs from the latter: It is independent of the stress response and of receptors needed for stress-induced ER-phagy, and while stress-induced ER-phagy is not discriminatory, constitutive ER-phagy has specific cargos. However, when constitutive ER-phagy fails, machinery specific to stress-induced ER-phagy can process its cargo. Moreover, constitutive ER-phagy is also not dependent on ER-stress or the unfolded protein response (UPR) stimulated by this stress, although its failure elicits UPR. Finally, constitutive ER-phagy and ERAD can partially process each other’s cargo upon failure. In summary, constitutive ER-phagy, which clears excess integral-membrane proteins from the ER during normal growth does not require nutritional or ER stress for its function.

## Introduction

The endoplasmic reticulum (ER) is the first stop for ~30% of all cellular proteins and as such is subject to a number of quality control (ER-QC) mechanisms [1]. The two destinations for excess or damaged cellular components are the proteasome and the lysosome. ER-associated degradation (ERAD) delivers misfolded proteins from the ER to the proteasome, especially during ER stress [2]. In contrast, ER fragments containing membrane and proteins can be delivered to the lysosome via micro- or macro-autophagy. In micro-autophagy, cellular components are enwrapped directly by the lysosomal membrane, while in macro-autophagy, they are first engulfed by the double-membrane autophagosome (AP), which later fuses with the lysosomal membrane. Macro-autophagy is induced by nutritional stress and requires a set of conserved autophagy-related proteins, Atgs [3, 4]. Nutritional stress can be induced in yeast either by nitrogen starvation, which signals through inactivation of the target of rapamycin complex 1 (TORC1), or by adding the drug rapamycin, which inhibits TORC1 directly [5]. In yeast, there is also a constitutive macro-autophagy pathway that during normal growth delivers proteins from the cytoplasm to the vacuole (the yeast lysosome), CVT [6]. However, knowledge about constitutive macro-autophagy in general is scarce in yeast and human cells.

We have previously reported that during normal growth, excess of integral-membrane proteins is shuttled from the ER for degradation in the vacuole via macro-autophagy. The idea that macro-ER-phagy clears this cargo is based on three findings: First, Atgs needed for AP formation (e.g., Atg1, Atg8, Atg9 and Atg2) are required for this process [7]. In addition, this process is regulated by a GTPase module that regulates AP biogenesis [8], which includes the Trs85-containing TRAPP III complex as an activator, the Ypt1 GTPase, and the Atg11 effector [9]. Second, the cargo passes through APs (shown by blocking a late autophagy step) [7]. Third, delivery of cargo through micro-autophagy was ruled out as micro-autophagy was intact in the cells defective for delivery of excess membrane proteins to the vacuole [7]. We identified two different proteins that upon overexpression serve as cargos for this pathway: GFP-Snc1-PEM (or DsRed-Snc1-PEM) and Snq2-yEGFP; Snc1 is a v-SNARE with a single trans-membrane domain (TMD) that mediates fusion of Golgi vesicles with the plasma membrane (PM), and Snq2 is a multi-TMD ABC transporter. Recently, we identified two endogenously-expressed ER resident proteins, Sec61 and Hmg1, as cargos for this pathway and showed that their delivery for degradation can be enhanced (2-4 fold) upon overexpression of a single integral-membrane protein [7]. Here, we wished to determine whether this ER-phagy pathway is constitutive.

To do that, we compared clearance of excess integral-membrane proteins during normal growth with nutritional stress-induced autophagy using several criteria. First, a group of core Atgs, like the yeast Atg8 and the mammalian LC3, plays a role both in nutritional stress-induced and constitutive autophagy, and one hallmark of nutritional stress-induced autophagy is an increase in the level of Atg8 [10]. Second, some Atgs, like Atg17 and Atg11, are specific to nutritional stress-induced and the constitutive autophagy pathway CVT, respectively [11, 12]. Finally, selective delivery of individual cellular components to the lysosome under stress requires cargo-specific Atg8/LC3 receptors that contain Atg8-interaction motif (AIM; LC3-interaction motif, LIR, in mammalian cells) and ensure effective packaging of the cellular components into Atg8/LC3-contianing APs [13, 14]. For the ER these selective receptors are Atg39 and Atg40 [15].

In addition, we explored here whether constitutive ER-phagy overlaps with ERAD that also occurs during normal growth. In ERAD, proteins that contain degradation signals termed degrons are ubiquitinated and degraded by the ubiquitin-proteasome system, UPS. Categorization of ERAD types depends on the localization of degrons in their substrates. Degron recognition by ERAD-L, ERAD-M, and ERAD-C, occurs in the ER lumen, ER membrane, and the cytoplasm, respectively. Doa10 and Hrd1 are conserved E3 ligases that target ERAD-C and ERAD-M/L substrates, respectively [16]. The two substrates for ER-phagy during normal growth, GFP-Snc1-PEM and Snq2-yEGFP [9], have large cytosolic domains and a single or multiple TMDs, respectively [17, 18]. Therefore, we explored the potential overlap between ER-phagy and ERAD-C or ERAD-M/L during normal cell growth.

Failure to clear misfolded proteins from the ER results in ER stress that in turn induces the unfolded protein response (UPR) [19]. This stress response halts protein translation, induces degradation of misfolded proteins by ERAD, and activates a signaling pathway for expression of molecular chaperones [20, 21]. In addition, ER stress induces micro-ER-phagy [22]. We showed that UPR induction is not necessary for clearance of excess integral-membrane proteins by ER-phagy, even though their accumulation in mutants defective in this clearance induces UPR [7]. The question was whether this UPR induction is required for clearance of excess integral-membrane proteins by ERAD.

Here, we show that clearance of excess integral-membrane proteins from the ER via autophagy during normal growth requires neither induction of general autophagy nor the two ER-phagy receptors Atg39 and Atg40. Because it happens during normal growth and is independent of both nutritional and ER stress, we define this process as constitutive ER-phagy and show that upon failure it can partially be rescued by stress-induced autophagy machinery and ERAD.

## Results

### Overexpression of GFP-Snc1-PEM does not induce general autophagy response

Under nutritional stress, cell growth is arrested, and macro-autophagy is induced. Because cells continue to grow when overexpressed integral-membrane proteins are shuttled to the vacuole for degradation through ER-phagy [7, 9], we hypothesized that this pathway is constitutive and does not require induction of general autophagy. Three different approaches were used to assess whether general autophagy response is induced in cells overexpressing GFP-Snc1-PEM: determining Atg8 protein level [23], assessing processing of the CVT cargo Ape1 [11], and measuring the alkaline phosphatase activity of Pho8Δ60 [24].

Wild-type and *atg11*Δ mutant cells were transformed with a 2μ plasmid for overexpression of GFP-Snc1-PEM (or with empty plasmid as a negative control). Snc1 is normally delivered to the PM and then cycles back through the Golgi. GFP-Snc1-PEM contains a modified TMD and two mutations that make it internalization defective [25]. Once GFP-Snc1-PEM reaches the PM, it stays there, and therefore any intracellular GFP signal is due to a block *en route* to the PM [9]. Experiments were performed with cells growing under normal conditions or under nitrogen starvation. The presence and level of GFP-Snc1-PEM was verified using immuno-blot and fluorescence microscopy analyses. Under normal growth conditions (SD+N), as we have previously shown, *atg11*Δ mutant cells accumulate 3.5-fold more GFP-Snc1-PEM when compared to wild-type cells (Figure 1A). As previously shown, GFP-Snc1-PEM (or GFP) does not accumulate in the vacuole neither in wild type nor in *atg11*Δ mutant cells, but it does accumulate in the majority of *atg11*Δ mutant cells outside the vacuole (Figure 1B) [7, 9]. Moreover, UPR induction was determined to confirm that this accumulation occurs in the ER and induces ER stress. In a comprehensive screen of the yeast gene deletion collection, *ATG* deletion mutants (including *atg11*Δ) were not identified among deletions that resulted in constitutive UPR upregulation [26]. Importantly, when GFP-Snc1-PEM is overexpressed, UPR is induced in *atg11*Δ, but not in wild type cells, indicating that the intracellular accumulation of GFP-Snc1-PEM occurs in the ER of the mutant cells ([9], and see Figure 2C). Under nitrogen starvation (SD-N), GFP-Snc1-PEM can be shuttled from the ER to the vacuole via the nutritional stress-induced autophagy, which is not dependent on Atg11 [27]. Indeed, the levels of GFP-Snc1-PEM in wild-type and *atg11*Δ mutant cells are similar and GFP fluorescence can be seen in vacuoles (outlined with the FM4-64 dye) of both strains (Figure 1A-B). The lower accumulation of GFP-Snc1-PEM in *atg11*Δ mutant cells upon nutritional stress is further discussed in the next section. Induction of general autophagy was assessed in these cells during normal growth (with nutritionally-stressed cells serving as positive control for the induction).

**Figure 1.**
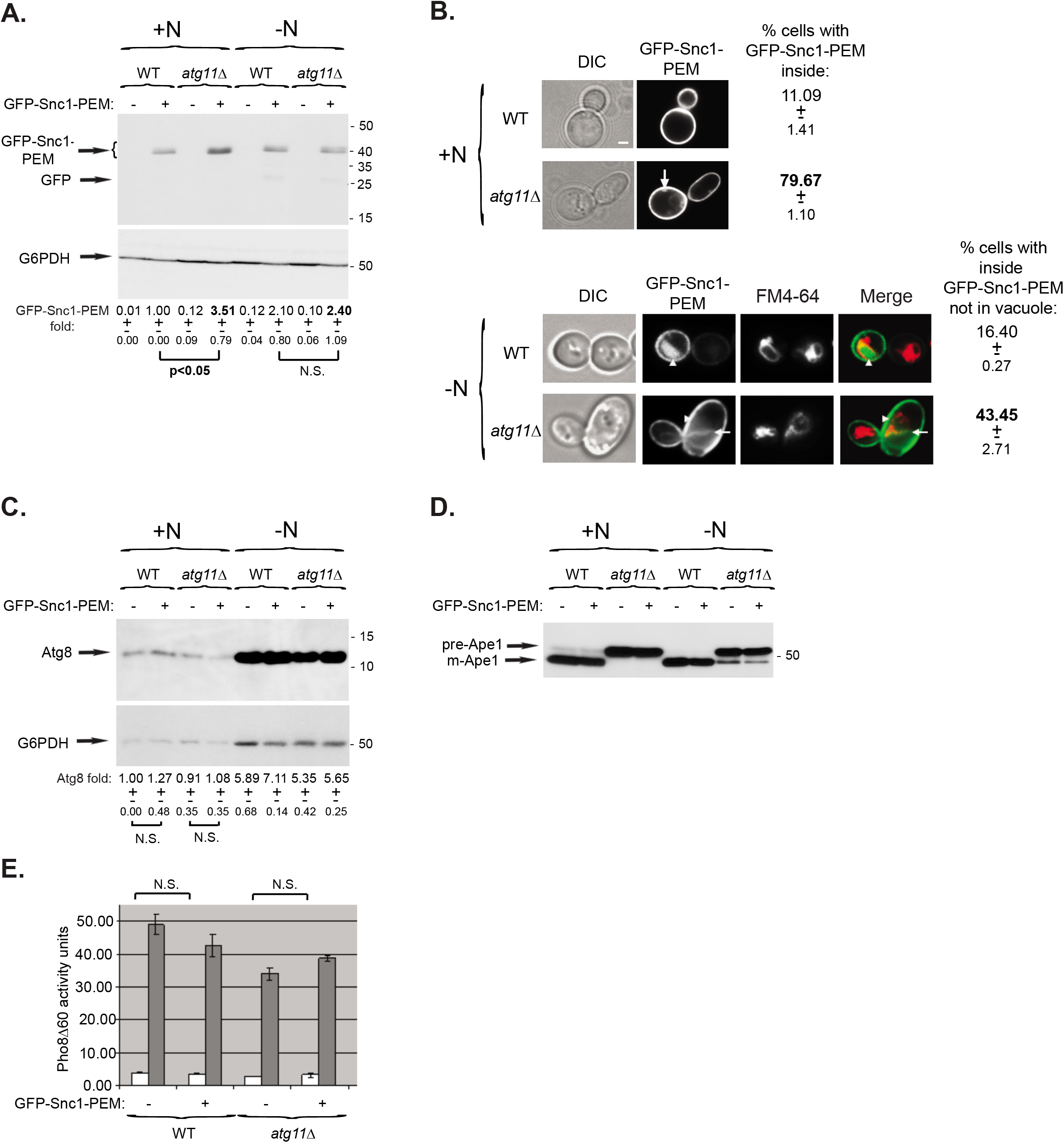
General autophagy is not induced during constitutive ER-phagy of an overexpressed membrane protein. **A-C.** Overexpression or intracellular accumulation of GFP-Snc1-PEM does not result in elevation of Atg8 protein level. **A.** *atg11*Δ mutant cells accumulate 3.5-fold more GFP-Snc1-PEM than WT cells during normal growth. WT and *atg11*Δ mutant cells were transformed with a 2μ plasmid for overexpression of GFP-Snc1-PEM (or empty plasmid as a negative control). Cells were grown either in SD+N medium (left), or in medium without N (right) for 6 hours. The level of GFP-Snc1-PEM in cell lysates was determined using anti-GFP antibodies and immuno-blot analysis (G6PDH was used as a loading control). Shown from top to bottom: growth medium, strain, plasmid, GFP blot, G6PDH blot, and quantification of the GFP-Snc1-PEM band: fold over wt, +/- STD, p-value. **B.** *atg11*Δ mutant cells accumulate intracellular GFP-Snc1-PEM during normal growth. Cells from panel A were visualized by live-cell fluorescence microscopy, except that FM4-64 was added to cells during nitrogen starvation. Shown from left to right: growth condition, strain, DIC (for cell contour), GFP, and % cells in which intracellular GFP-Snc1-PEM accumulates in the cytoplasm (not in vacuole); +/-, STD, p-value. GFP-Snc1-PEM and GFP accumulates in the cytoplasm of the majority of *atg11*Δ mutant cells during normal growth (top), and ~2-fold less under starvation (bottom); (p-value <0.01). Under nitrogen starvation (-N), GFP-Snc1-PEM accumulates in the vacuoles (outlined with FM4-64) of both WT and mutant cells. >100 cells were visualized per panel; arrows and arrowheads point to GFP-Snc1-PEM in the cytoplasm and the vacuole, respectively; size bar, 1μ. **C.** The level of Atg8 is not increased due to overexpression of GFP-Snc1-PEM during normal growth, but is increased by 5-7-fold during N starvation regardless of GFP-Snc1-PEM overexpression. Cell lysates from panel A were tested by immuno-blot analysis using anti-Atg8 antibodies. Shown from top to bottom: Atg8 blot, G6PDH blot (loading control), quantification of fold Atg8 compared to the level in WT cells (+/-, STD), and p-values >0.05 showing no significant difference (N.S.) with and without Snc1 overexpression. **D.** Ape1 processing is not induced in *atg11*Δ mutant cells overexpressing GFP-Snc1-PEM. Cell lysates from panel A were analyzed by immuno-blot analysis for processing of prApe1 to mApe1 using anti-Ape1 antibodies. In WT cells, Ape1 is mostly processed during normal growth and fully processed under stress (-N). In *atg11*Δ mutant cells, Ape1 is partially processed under stress (-N), but not during normal growth regardless if the cells express GFP-Snc1-PEM. **E.** Pho8Δ60 phosphatase activity is not induced in WT or *atg11*Δ mutant cells expressing GFP-Snc1-PEM during normal growth. Pho8Δ60 phosphatase activity was measured in lysates of cells from panel A grown in SD+N (while bars); the activity is elevated in all cells under stress (gray bars, SD-N) regardless if they express GFP-Snc1-PEM. Error bars, STD. Results in this figure represent 3 independent experiments.

**Figure 2.**
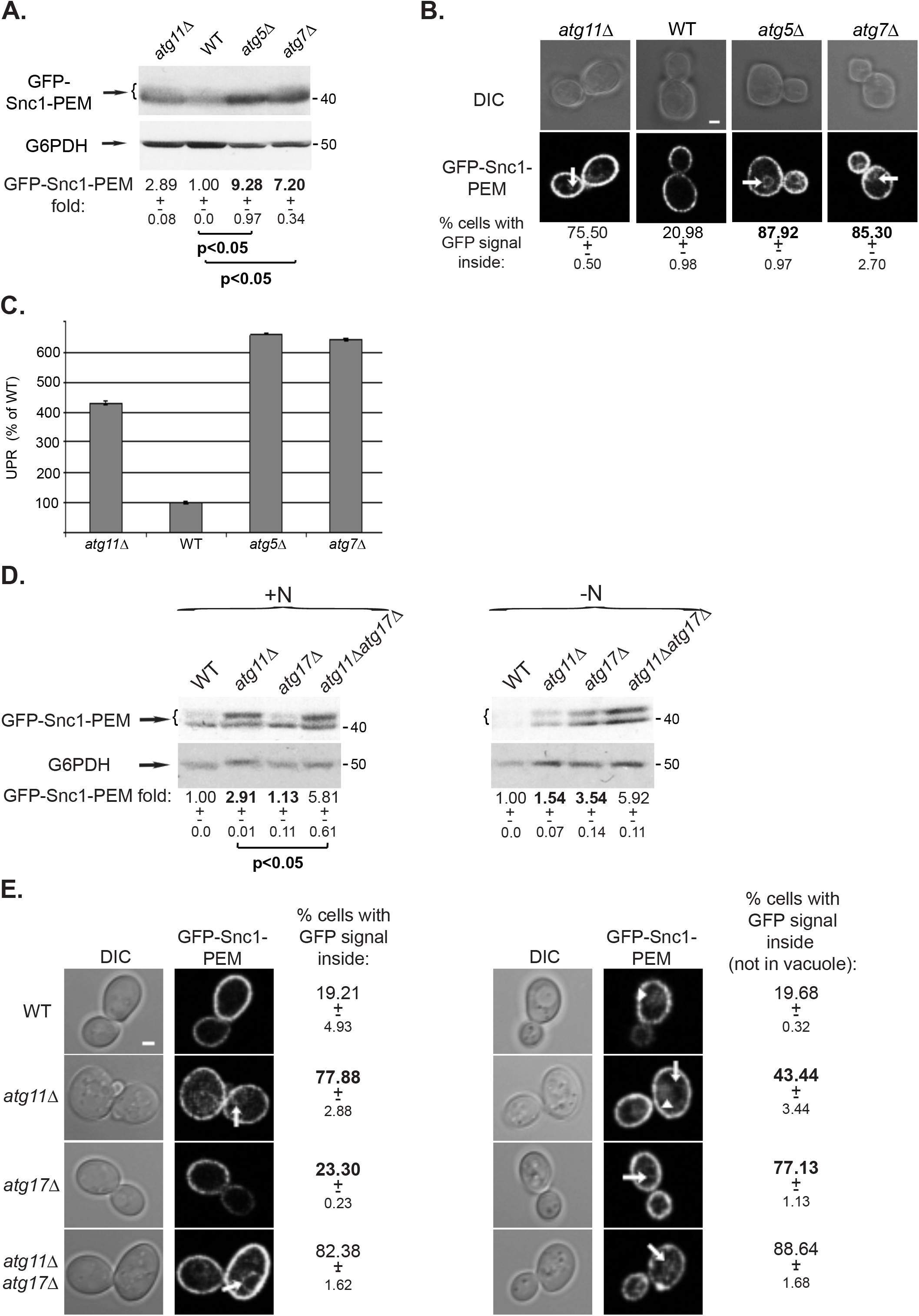
Contribution of core Atgs to autophagy of GFP-Snc1-PEM. **A-C.** Atg5 and Atg7 are required for constitutive ER-phagy. **A.** *atg5*Δ and *atg7*Δ mutant cells accumulate GFP-Snc1-PEM. WT and mutant cells (*atg11*Δ as a reference) transformed with a 2μ plasmid for overexpression of GFP-Snc1-PEM, were grown under normal growth conditions (SD+N). The level of GFP-Snc1-PEM in their lysates was tested by immuno-blot analysis using anti-GFP antibodies. Shown from top to bottom: strain, GFP blot, G6PDH blot (loading control), fold Snc1-PEM-GFP over the WT level, +/- STD, p-value. **B.** GFP-Snc1-PEM is accumulated inside of the majority of *atg5*Δ and *atg7*Δ mutant cells. Cells from panel A were visualized by live-cell fluorescence microscopy. Shown from top to bottom: strain, DIC, GFP, and percent cells with intracellular GFP-Snc1-PEM +/- STD. >100 cells were visualized per panel; arrows point to GFP-Snc1-PEM in the cytoplasm; scale bar, 1μ. **C.** UPR is induced in *atg5*Δ and *atg7*Δ mutant cells that overexpress GFP-Snc1-PEM. Cells from panel A were transformed with a second plasmid for expression of ßgal under UPR response elements and the induction of the ßgal activity was determined in cell lysates. Bar graph shows % of UPR compared to WT (100%); error bars, STD. Like in *atg11*Δ mutant cells, UPR is induced in *atg5*Δ and *atg7*Δ mutant cells (~6.5-fold). **D-E.** Atg11 and Atg17 play major roles in autophagy of GFP-Snc1-PEM during normal growth and under stress (-N), respectively. **D.** Most of the GFP-Snc1-PEM that accumulates in *atg11*Δ mutant cells during normal growth is degraded under stress (-N) in an Atg17-dependent manner. Wild-type, *atg11*Δ, *atg17*Δ and *atg11*Δ *atg17*Δ mutant cells transformed with a 2μ plasmid for overexpression of GFP-Snc1-PEM were grown under normal growth conditions (SD+N, left) or under starvation (SD-N, right). The level of GFP-Snc1-PEM in cell lysates was determined using anti-GFP antibodies and immuno-blot analysis (G6PDH was used as a loading control). Shown from top to bottom: growth medium, strain, GFP blot, G6PDH blot, and quantification of the GFP-Snc1-PEM band: fold over wt, +/- STD, p-value. **E.** Under stress (-N), the portion of *atg11*Δ that accumulate GFP-Snc1-PEM is decreased in an Atg17-dependent manner. Cells from panel D were visualized by live-cell fluorescence microscopy. Shown from left to right: DIC (for cell contour), GFP, and % cells in which intracellular GFP-Snc1-PEM accumulates in the cytoplasm (not in vacuole); +/-, STD. GFP-Snc1-PEM accumulates in the cytoplasm of the majority of *atg11*Δ mutant cells during normal growth (left), and ~2-fold less under starvation (right) (p-value <0.02). Under nitrogen starvation(-N), GFP-Snc1-PEM accumulates in the vacuoles of WT and *atg11*Δ, but not *atg17*Δ or *atg11*Δ *atg17*Δ, mutant cells. >100 cells were visualized per panel; arrows and arrowheads point to GFP-Snc1-PEM in the cytoplasm and the vacuole, respectively; size bar, 1μ. Results in this figure represent 3 independent experiments.

First, as expected, the level of Atg8 protein is induced ~5-7-fold under nitrogen starvation in both wild type and *atg11*Δ mutant cells regardless of the expression of GFP-Snc1-PEM (Figure 1C, right). In contrast, the level of Atg8 is not elevated in wild-type and *atg11*Δ mutant cells expressing GFP-Snc1-PEM, when compared to cells that do not overexpress this membrane protein (Figure 1C, left). Thus, neither sending GFP-Snc1-PEM for degradation through autophagy (in wild-type cells) nor its accumulation (in *atg11*Δ mutant cells) results in elevation of Atg8 protein level.

Second, Ape1 processing from prApe1 to mApe1 is mostly dependent on Atg11 [27], but, there are differences in this processing in both wild-type and *atg11*Δ mutant cells during normal growth and under starvation. In wild-type cells, while the majority of Ape1 is processed during normal growth, it is fully processed under starvation, when general autophagy is induced. In *atg11*Δ mutant cells, Ape1 processing is completely blocked under normal growth, but some of it is processed under starvation. Importantly, overexpression of GFP-Snc1-PEM does not cause a change in Ape1 processing during normal growth in wild-type or *atg11*Δ mutant cells (Figure 1D), indicating that general autophagy was not induced.

Third, under nitrogen starvation, alkaline phosphatase activity of Pho8Δ60 is increased in both wild type and *atg11*Δ mutant cells regardless of the presence of GFP-Snc1-PEM ([9] and Figure 1E, gray bars). Importantly, under normal growth conditions, Pho8Δ60 activity is not increased neither in wild type cells, not in *atg11*Δ mutant cells expressing GFP-Snc1-PEM (Figure 1E, white bars). Thus, general autophagy is not induced when ER-phagy of a membrane protein occurs (in wild type cells) or blocked (in *atg11*Δ mutant cells).

At least for one type of alternative macro-autophagy of mitochondria that occurs during maturation of mouse erythroids, Atg5 and Atg7, which mediate Atg8 lipidation, are not required [28]. Because Atg8 level does not increase in the ER-phagy characterized here, we tested the requirement of Atg5 and Atg7 for this process. When GFP-Snc1-PEM was overexpressed in cells deleted for *ATG5* or *ATG7*, ER-phagy was blocked as evident from intracellular accumulation of GFP-Snc1-PEM (similar to *atg11*Δ), its increased level (7-9-fold), and an increase in the UPR response in these mutant cells (>6-fold) (Figure 2A-C).

Together, our results show that even though the Atg machinery, including Atg8 and its lipidation, is required for ER-phagy of overexpressed integral-membrane proteins, general autophagy is not induced while it functions. This is based on the facts that cells continue to grow and lack two hallmarks of nutritional stress-induced autophagy, increase in Atg8 protein and Pho8Δ60 activity levels. Hereafter, we term this process constitutive ER-phagy, with the caveat that, as we showed previously [7], shuttling of endogenously expressed ER membrane proteins through it can be further induced when a single integral-membrane protein is overexpressed.

### Relationship between constitutive and nutritional stress-induced ER-phagy

#### Shuttling of constitutive ER-phagy cargo for degradation under nutritional stress

In yeast, two Atgs are specific to constitutive versus nutritional stress-induced autophagy, Atg11 and Atg17, respectively [4]. What happens to the cargo of constitutive ER-phagy that accumulates in mutant cells defective in this process during normal growth (e.g., *atg11*Δ) after induction of nutritional stress? While during normal growth delivery of overexpressed GFP-Snc1-PEM to the vacuole depends mostly on Atg11 and not on Atg17, under nutritional stress this cargo is delivered for degradation to the vacuole mainly via an Atg17-dependent pathway. Under both growth conditions, the double mutant *atg11*Δ *atg17*Δ exhibits the most severe block. This can be seen both by immuno-blot and fluorescence microscopy analyses (Figure 2 D-E). Additive effects of *atg11*Δ and *atg17*Δ have been previously reported in other autophagy types [11] and we reported this for GFP-Snc1-PEM accumulation during normal growth [7]. This ~2-fold increase suggests that during normal growth in the absence of Atg11, Atg17-dependent autophagy, that usually functions under stress, can clear about 50% of the accumulated cargo.

Interestingly, stress (-N) partially relieves the *atg11*Δ block because the accumulation of GFP-Snc1-PEM goes down from ~3- to 1.5-fold, and the percent of cells accumulating intracellular GFP-Snc1-PEM goes down from ~80% to 43%. Thus, ~50% of GFP-Snc1-PEM that accumulates in *atg11*Δ mutant cells during normal growth can be degraded under nutritional stress and Atg17 is required for that, because the double mutant *atg11*Δ *atg17*Δ accumulates more GFP-Snc1-PEM in more cells than the single mutants. Thus, under nutritional stress, the constitutive ER-phagy cargo behaves like all the other ER cargos, and as shown below, is also dependent on nutritional stress-induced selective ER-phagy receptors.

#### Receptors for nutritional stress-induced ER-phagy and their cargo

ER-phagy that occurs under nutritional stress requires the Atg8-receptors Atg39 or Atg40, which are ER resident proteins that ensure engulfment of ER fragments by APs. While single deletions exhibit partial defects in this nutritional stress-induced ER-phagy, *atg39*Δ *atg40*Δ double deletion results in a complete block of this process [15]. We wished to determine whether Atg39 and Atg40 play a role in constitutive ER-phagy of an overexpressed integral-membrane protein and whether they are sorted for delivery to the vacuole through constitutive ER-phagy.

The ability of *atg39*Δ *atg40*Δ double mutant cells to send overexpressed GFP-Snc1-PEM for degradation in the vacuole through ER-phagy was determined both during normal growth and under nutritional stress. Under normal growth conditions (SD+N), GFP-Snc1-PEM level is not increased, it does not accumulate inside cells, and UPR is not increased (Figure 3A-C) in *atg39*Δ *atg40*Δ double mutant cells. Moreover, combining the *atg39*Δ *atg40*Δ deletions with *atg11*Δ or *atg17*Δ does not result in more severe phenotypes than those of the single *atg11*Δ or *atg17*Δ mutant cells (Figure 3A-C). In contrast, as expected, under starvation (SD-N), *atg39*Δ *atg40*Δ mutant cells exhibit defects in shuttling GFP-Snc1-PEM to the vacuole for degradation as attested by the increase of GFP-Snc1-PEM level (3-fold) and its accumulation inside cells (Figure 3A-B, right). Interestingly, under starvation more GFP-Snc1-PEM accumulates in *atg17*Δ than in *atg39*Δ *atg40*Δ mutant cells (~4- and 3-fold, respectively, Figure 3A, -N). This suggests that during nutritional stress, some ER cargo can be delivered to the vacuole in an Atg17-dependent way either via other receptors or in bulk. Under starvation, combining *atg39*Δ *atg40*Δ deletions with *atg17*Δ or *atg11*Δ does not result in additive phenotypes, suggesting that they function in the same pathway. These results show that while GFP-Snc1-PEM is shuttled to the vacuole through Atg39/Atg40-mediated selective ER-phagy during nutritional stress, these two receptors are not important for constitutive ER-phagy during normal growth.

**Figure 3.**
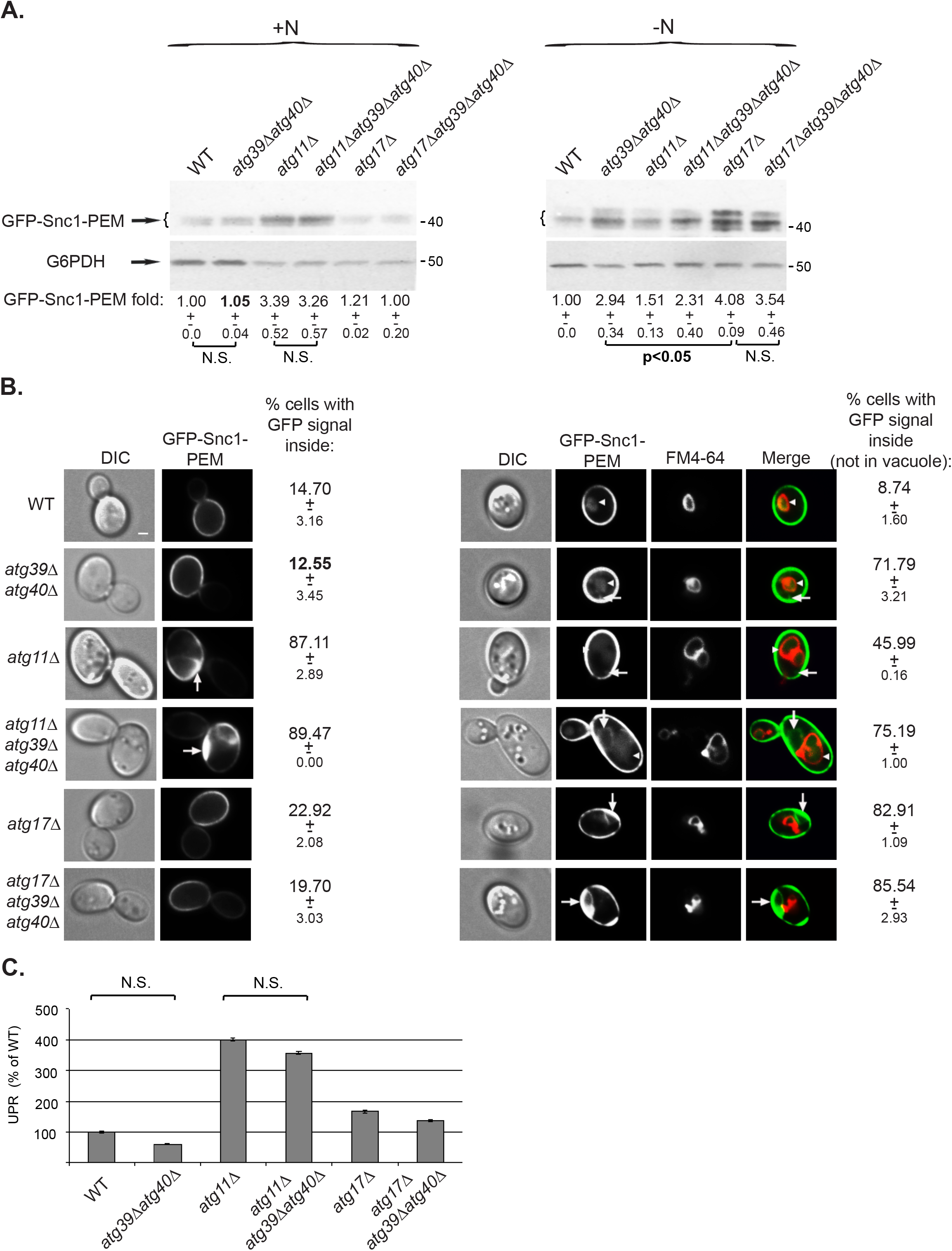
The AIM receptors Atg39 and Atg40 are not required for constitutive ER-phagy. **A.** *atg39*Δ *atg40*Δ double deletion, alone or in combination with *atg11*Δ or *atg17*Δ, does not cause increase in GFP-Snc1-PEM protein level during normal growth. WT and indicated mutant cells transformed with a 2μ plasmid for overexpression of GFP-Snc1-PEM, were grown in normal (SD+N, left) or starvation medium (SD-N, right). The level of Snc1-PEM-GFP in cell lysates was tested by immuno-blot analysis using anti-GFP antibodies. Under starvation, *atg39*Δ *atg40*Δ causes increase in GFP-Snc1-PEM level, showing that these receptors are important for starvation-induced ER-phagy. Shown from top to bottom: strain, GFP blot, G6PDH blot (loading control), fold GFP-Snc1-PEM over the WT level; +/- STD, p-value. **B.** *atg39*Δ *atg40*Δ double deletion, alone or in combination with *atg1*Δ or *atg17*Δ, does not result in intracellular accumulation of GFP-Snc1-PEM during normal growth. Cells from panel A were grown in SD+N (left), or in SD-N (stained by FM4-64 to label the vacuolar membrane; right), and were visualized by fluorescence live-cell microscopy. Under starvation, *atg39*Δ *atg40*Δ causes increase in intracellular Snc1-PEM-GFP that accumulates outside the vacuole, showing that these receptors are important for starvation-induced ER-phagy. Shown from left to right: strain, DIC, GFP, (FM4-64, merge, for SD-N), and percent cells with intracellular GFP-Snc1-PEM (not in vacuole) +/- STD. >100 cells were visualized per panel; arrows and arrowheads point to Snc1-PEM-GFP in the cytoplasm and the vacuole, respectively; size bar, 1μ. **C.** UPR is not induced in *atg39*Δ *atg40*Δ mutant cells that overexpress GFP-Snc1-PEM during normal growth. Induction of the UPR response was determined as explained for Figure 2C. Bar graph shows % of UPR compared to WT (100%). While UPR is induced in *atg11*Δ mutant cells, it is not induced in *atg39*Δ *atg40*Δ double deletion mutant cells; error bars, STD. Results in this figure represent 3 independent experiments.

Under nutritional stress, Atg39 and Atg40 are delivered to the vacuole for degradation [15]. Since they are both ER integral-membrane proteins, we wished to determine whether these two proteins can act as cargos of the constitutive ER-phagy. As was previously shown [15], when Atg39 and Atg40 were tagged with HA at their C-termini they retain their functionality, because cells expressing them as the only copy of Atg39 and Atg40 can still degrade their respective ER cargos, Nop1 and Rtn1, under nutritional stress (Figure S1). We followed the levels of endogenously-expressed HA-tagged Atg39 and Atg40 during normal growth and under nutritional stress, and during normal growth of cells overexpressing GFP-Snc1-PEM. During normal growth (no rapamycin), Atg39 and Atg40 are stable as seen by their similar levels in wild type and cells deleted for vacuolar proteases (*pep4*Δ *prb1*Δ),indicating that they were not delivered to the vacuole for degradation (Figure S2, top). As expected, under stress (+rapamycin), Atg39 and Atg40 are delivered to the vacuole for degradation, because their levels are higher in cells deleted for *pep4*Δ *prb1*Δ when compared to wild-type cells (Figure S2, bottom). Interestingly, in *ypt1-1* mutant cells, the levels of Atg39 and Atg40 are lower when compared to wild-type cells during normal growth and under stress. Because this decrease can be partially suppressed by depletion of vacuolar proteases (Figure S2), Atg39 and Atg40 can probably be delivered to the vacuole for degradation in a Ypt1-independent manner.

Importantly, during normal growth of cells overexpressing GFP-Snc1-PEM, while GFP-Snc1-PEM accumulates in *ypt1-1* mutant cells, neither Atg39 nor Atg40 proteins accumulate. In fact, not only Atg39 and Atg40 do not accumulate, their level is lower than that of the wild type as observed in cells that do not overexpress GFP-Snc1-PEM (above). Moreover, while in wild-type cells depleted for vacuolar proteases (*pep4*Δ *prb1*Δ) GFP-Snc1-PEM accumulates, indicating it was shuttled to the vacuole for degradation, the levels of Atg39 and Atg40 do not change (Figure 4A-B).

**Figure 4.**
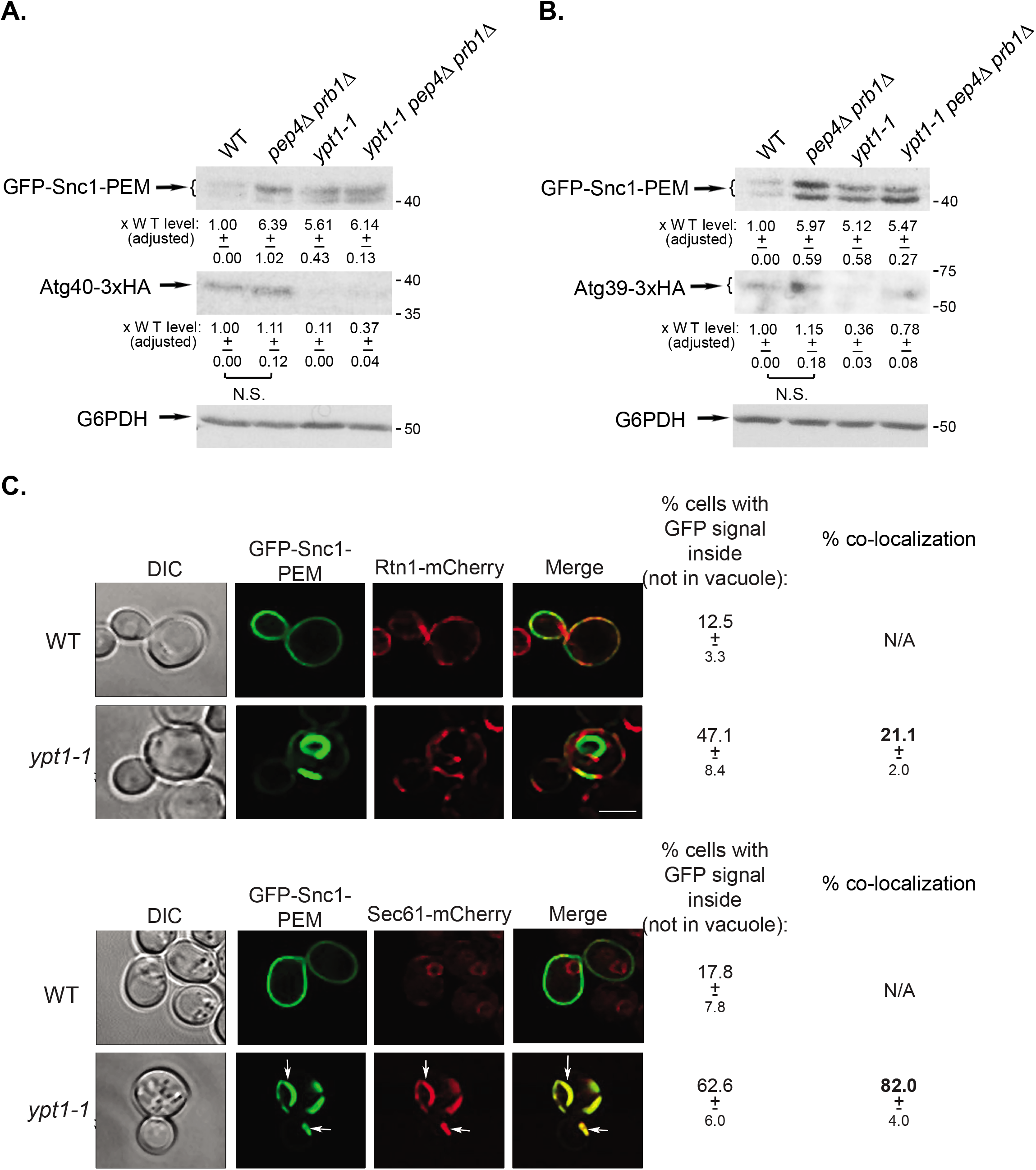
Neither Atg39 and Atg40 nor the Atg40 cargo Rtn1 are shuttled through constitutive ER-phagy. **A-B.** The ER membrane proteins Atg39 and Atg40 do not accumulate in *ypt1-1* mutant cells. Endogenous Atg39 and Atg40 were tagged with 3xHA at their C-termini. The cells were transformed with a 2μ plasmid for overexpression of GFP-Snc1-PEM and grown at 26°C (a permissive temperature for *ypt1-1* mutant cells growth in SD, [9]). The level of GFP-Snc1-PEM and Atg40 (**A**) or Atg39 (**B**) in cell lysates was determined using anti-GFP and anti-HA antibodies and immuno-blot analysis (G6PDH was used as a loading control). Shown from top to bottom: strain, GFP blot, HA blot, quantification of the HA band: fold over wt, +/- STD, p-value, and G6PDH blot. **C.** The ER protein Rtn1 does not accumulate inside *ypt1-1* mutant cells and does not co-localize with GFP-Snc1-PEM. Endogenous Rtn1 was tagged at its C-terminus with mCherry. The cells were transformed with a 2μ plasmid for overexpression of GFP-Snc1-PEM and visualized by live-cell microscopy (top). Cells expressing Sec61-mCherry were used as a positive control for co-localization with GFP-Snc1-PEM (bottom). Shown from left to right: strain, DIC, GFP, mCherry, Merge, percent cells with intracellular GFP-Snc1-PEM (not in vacuole) +/- STD, and % co-localization of GFP-Snc1-PEM with the ER protein, Rtn1 or Sec61 in *ypt1-1* mutant cells. Four-fold less Rtn1 overlaps with GFP-Snc1-PEM in *ypt1-1* mutant cells than Sec61 (p-value<0.0001). >150 cells were visualized per panel; arrows point to sites where GFP-Snc1-PEM and the mCherry tagged ER proteins, Sec61 or Rtn1, co-localize; size bar, 5μ. Results in this figure represent 3 independent experiments.

Together, these results show that the selective Atg8-receptors, Atg39 and Atg40, which play a role in nutritional stress-induced ER-phagy, are not required for constitutive ER-phagy nor can they serve as cargos for this pathway.

If the ER-phagy receptors do not serve as cargos for constitutive-ER-phagy, what about their cargos? We tested it for Rtn1, an ER integral-protein whose delivery to the vacuole under nutritional stress depends on Atg40 [15]. Endogenous Rtn1 was tagged at its C-terminus with mCherry in wild-type, *atg40*Δ, and *ypt1-1* mutant cells. During normal growth, Rtn1-mCherry resides on the ER under the PM [15]. We confirmed that Rnt1-mCherry is an Atg40 cargo because under stress (induced by rapamycin), it is shuttled to the vacuole in wild-type, but not *atg40*Δ mutant cells (Figure S3). Cells expressing Rtn1-mCherry, and Sec61-mCherry for comparison, were transformed with a 2μ plasmid for overexpression of GFP-Snc1-PEM and visualized by live-cell microscopy. In wild-type cells, while Sec61 is present on the ER around the nucleus, Rtn1 is present on the ER under the PM (Figure 4C). We have previously shown that Sec61-mCherry accumulates together with GFP-Snc1-PEM in the cytoplasm of ~60% of *ypt1-1* mutant cells ([7], and Figure 4C, bottom, 82% co-localization). In contrast, in *ypt1-1* mutant cells expressing Rtn1-mCherry, GFP-Snc1-PEM accumulates in ~50% of the cells, but its co-localization with Rtn1-mCherry is 4-fold lower (Figure 4C, top, 21% co-localization). Thus, the Atg40 cargo Rtn1 does not accumulate in *ypt1-1* mutant cells together with GFP-Snc1-PEM.

In summary, whereas the Atg40 receptor and its ER cargo Rtn1 are shuttled to the vacuole through nutritional stress-induced ER-phagy, neither serves as a cargo for constitutive ER-phagy.

### Relationship between constitutive ER-phagy and ERAD

To understand whether constitutive ER-phagy and ERAD can supplement each other’s function in quality control of the ER, we used cargos specific to each pathway and mutants defective in each pathway: For constitutive ER-phagy we used *atg11*Δ, *trs85*Δ, *atg5*Δ and *atg9*Δ, for ERAD-C, we used *doa10*Δ, and for ERAD-M/L we used *hrd1*Δ.

#### Constitutive ER-phagy cargo and ERAD-C

Doa10 is a ubiquitin ligase that resides in the ER, recognizes cytosolic domains of misfolded ER membrane proteins, and facilitates their degradation via ERAD-C [29, 30]. Accumulation of the constitutive ER-phagy cargo GFP-Snc1-PEM in *doa10*Δ mutant cells was compared to that of *atg11*Δ or *trs85*Δ mutant cells under normal growth conditions (SD+N) using three parameters: increase in the level of GFP-Snc1-PEM, its intracellular accumulation and increase in UPR. Unlike *atg11*Δ and *trs85Δ, doa10*Δ mutant cells do not exhibit GFP-Snc1-PEM increase, its intracellular accumulation and increase in UPR (Figure 5). However, when *doa10*Δ is combined with *atg11*Δ or *trs85*Δ, the double deletion mutant cells exhibit a more severe phenotype and accumulate 35% or 50% more GFP-Snc1-PEM than the single deletions, respectively (Figure 5A and D). This increase was only manifested by the increase in GFP-Snc1-PEM protein level and not by the percent of cells accumulating intracellular GFP-Snc1-PEM or by the level of UPR induction (Figure 5B-C and E-F). This suggests that in mutant cells defective in constitutive ER-phagy, e.g., *atg11*Δ or *trs85*Δ, Doa10-mediated ERAD can partially help in clearance of GFP-Snc1-PEM.

**Figure 5.**
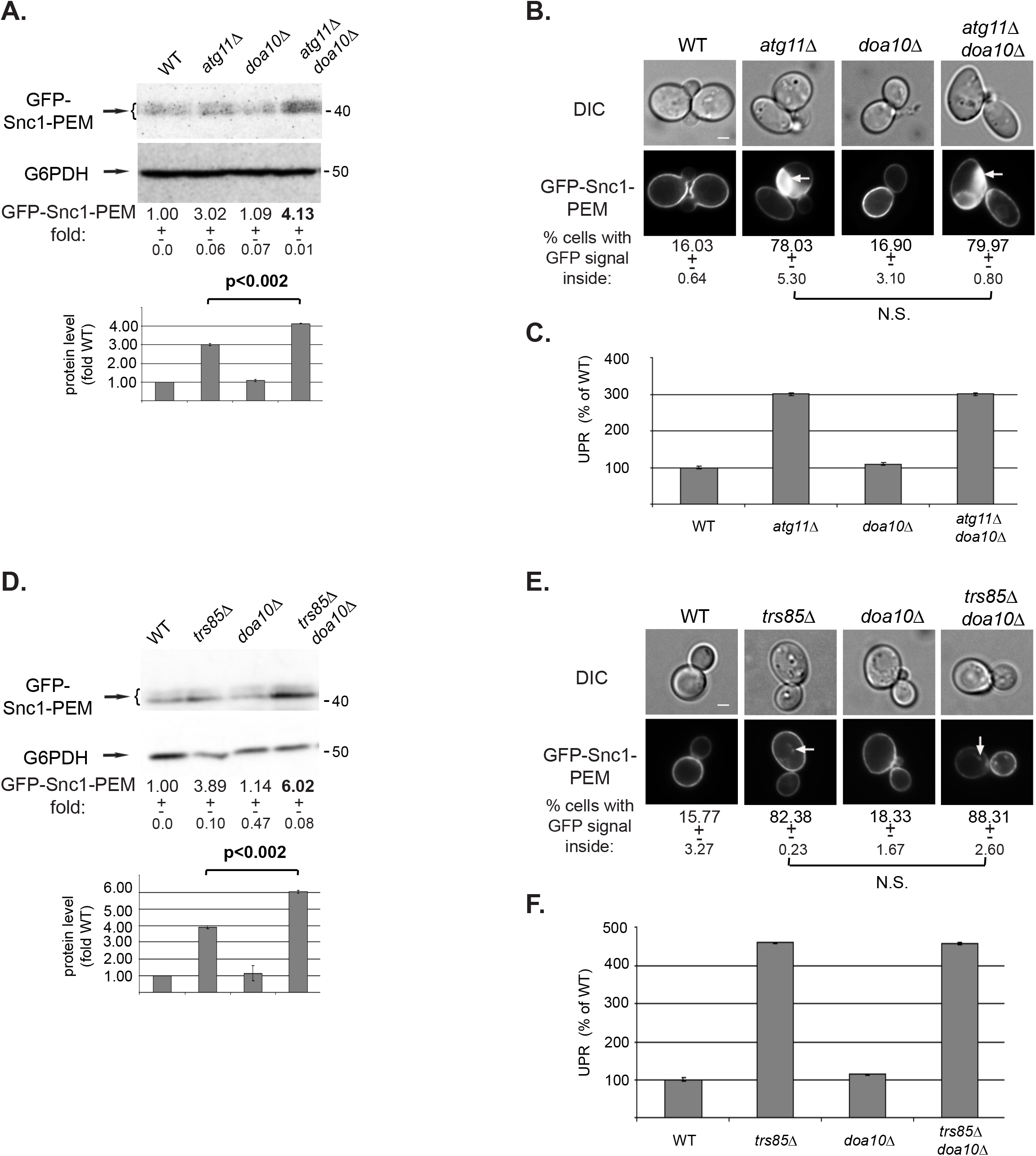
While deletion of *DOA10* alone does not result in GFP-Snc1-PEM accumulation, it does enhance the defect of *atg11*Δ and *trs85*Δ. **A-C.** Effect of *doa10*Δ alone and in combination with *atg11*Δ. **D-F.** Effect of *doa10*Δ alone and in combination with *trs85*Δ. WT and indicated mutant cells overexpressing GFP-Snc1-PEM were grown in normal growth medium (SD+N). The level of GFP-Snc1-PEM in cell lysates was determined by immuno-blot analysis using anti-GFP antibodies (**A, D**); intracellular accumulation of GFP-Snc1-PEM was determined by live-cell fluorescence microscopy (**B, E**); and UPR induction was analyzed (**C, F**). Results are presented as in Figure 2, except that the increase in Snc1-PEM-GFP protein level is shown as a bar graph under the blots, with p-value. A 35% and 50% increase in the level of GFP-Snc1-PEM is observed when *doa10*Δ is combined with *atg11*Δ and *trs85*Δ, respectively. Results in this figure represent 3 independent experiments.

To determine whether Doa10-mediated ERAD-C can partially clear GFP-Snc1-PEM that accumulate in cells deleted for Atgs required for general autophagy, we tested Atg5 and Atg9. Similar to *atg11*Δ and *trs85*Δ mutant cells, *doa10*Δ *atg5*Δ double mutant cells accumulated more GFP-Snc1-PEM than the single *atg5*Δ mutant cells (35% increase) (Figure S5A-B). Interestingly, *doa10*Δ does not exacerbate the ER-phagy phenotype of *atg9*Δ mutant cells (Figure 6A-C). We have previously shown that unlike *atg11*Δ and *trs85*Δ, which accumulate ER fragments decorated with GFP-Snc1-PEM, *atg9*Δ elicits in an earlier block in exit of GFP-Snc1-PEM from the ER before the formation of ER membranes containing GFP-Snc1-PEM. In addition, unlike *atg11*Δ and *trs85*Δ, UPR is not induced in *atg9*Δ mutant cells [7]. Thus, the inability of Doa10 to contribute to GFP-Snc1-PEM clearance in *atg9*Δ mutant cells could be because the ability of Doa10 to clear this cargo requires UPR induction or because Doa10 does not recognize the cargo accumulating in *atg9*Δ mutant cells.

**Figure 6.**
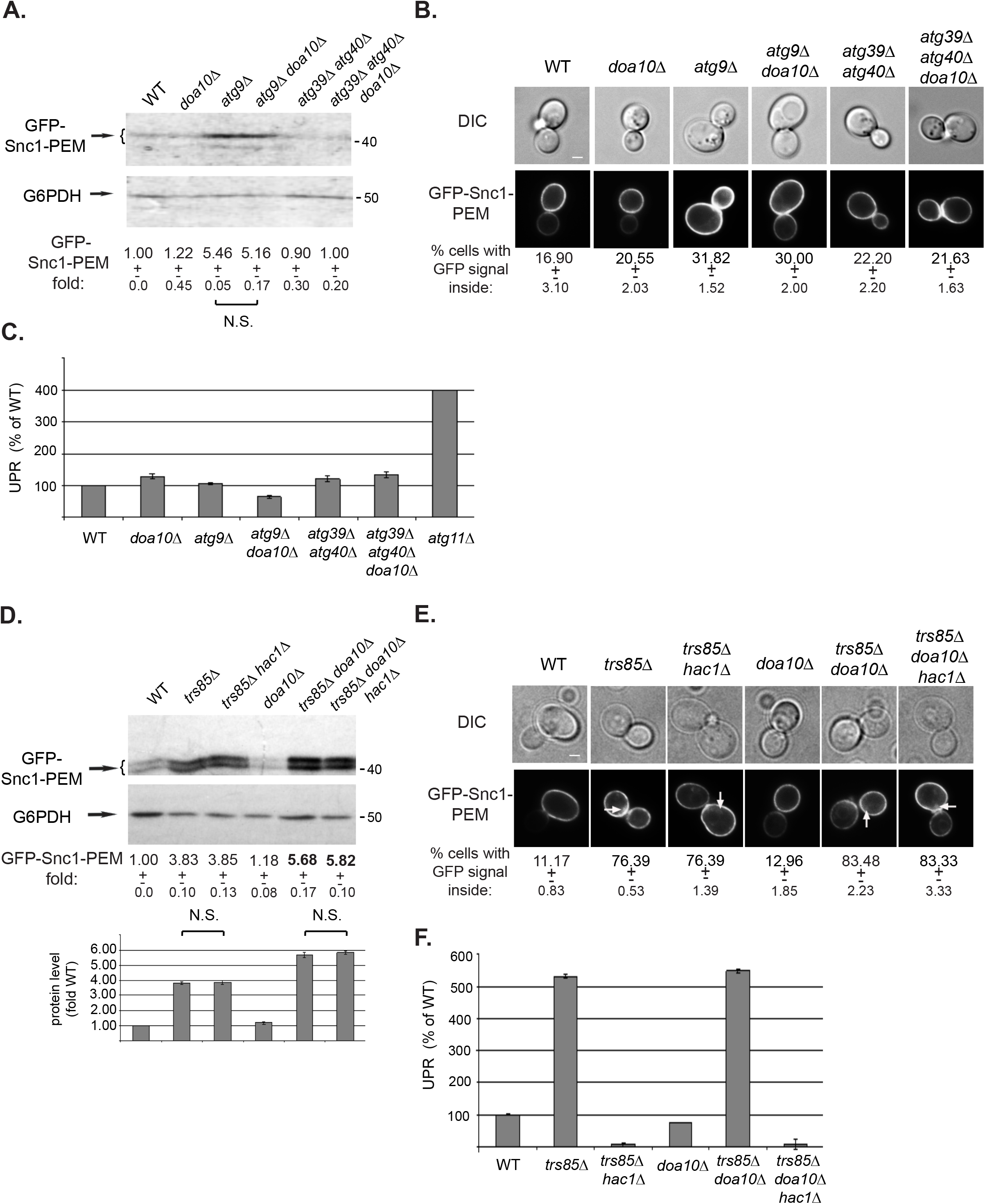
*doa10*Δ does not increase the constitutive ER-phagy defect of *atg9*Δ mutant cells, and its effect on *trs85*Δ phenotype is not dependent on UPR. **A-C.** Effect of *doa10*Δ alone and in combination with *atg9*Δ and *atg39Δ atg40Δ.* WT and indicated mutant cells overexpressing GFP-Snc1-PEM were grown in normal growth medium (SD+N). The level of GFP-Snc1-PEM in cell lysates was determined by immuno-blot analysis using anti-GFP antibodies (**A**); intracellular accumulation of GFP-Snc1-PEM was determined by live-cell fluorescence microscopy (**B**); and UPR induction was analyzed (**C**). Results are presented as in Figure 2. *doa10*Δ has no effect on GFP-Snc1-PEM accumulation in combination with *atg9*Δ, which exhibits a constitutive ER-phagy phenotype, or *atg39*Δ *atg40*Δ double deletion, which does not exhibit a constitutive ER-phagy phenotype. **D-F.** Abrogation of the UPR response by *hac1*Δ does not affect the constitutive ER-phagy phenotype of *trs85*Δ, or its enhancement in *trs85*Δ *doa10*Δ mutant cells. WT and indicated mutant cells overexpressing GFP-Snc1-PEM were grown in normal growth medium (SD+N). The level of GFP-Snc1-PEM in cell lysates was determined by immuno-blot analysis using anti-GFP antibodies (**D**); intracellular accumulation of GFP-Snc1-PEM was determined by live-cell fluorescence microscopy (**E**); and UPR induction were analyzed as explained for Figure 2C (**F**). The 50% increase in the GFP-Snc1-PEM accumulation phenotype in *trs85Δ doa10*Δ double mutant cells is affected by *hac1*Δ (panel D). Results are presented as in Figure 2 and represent 3 independent experiments.

To test the requirement of UPR for the contribution of Doa10 in clearing GFP-Snc1-PEM, UPR was blocked using *hac1*Δ [19]. We have previously shown that UPR is induced in most mutant cells defective in clearance of GFP-Snc1-PEM, except in *atg9*Δ, even though UPR is not required for constitutive ER-phagy to happen [7]. As expected, combining *hac1*Δ with *trs85*Δ did not affect the GFP-Snc1-PEM accumulation phenotype of *trs85*Δ mutant cells. Likewise, combining *hac1*Δ with *trs85Δ doa10*Δ did not affect the GFP-Snc1-PEM accumulation phenotype of the double deletion (Figure 6D-F). Thus, UPR induction is not necessary for Doa10 to help with the clearance of GFP-Snc1-PEM from the ER in the absence of Trs85. Therefore, the inability of Doa10 to help with GFP-Snc1-PEM clearance in *atg9*Δ mutant cells is not due to the lack of UPR induction. Rather, the evidence points to the inability of Doa10 to recognize the cargo in *atg9*Δ mutant cells.

While like Doa10, Atg39 and Atg40 are not required for constitutive ER-phagy (see above), if they played a role in a constitutive ER housekeeping process, an interplay between them and Doa10 would be expected. In this case, deleting all three genes would result in a constitutive ER-phagy phenotype. However, the triple deletion mutant *atg39*Δ *atg40*Δ *doa10* does not show any defect in GFP-Snc1-PEM accumulation and/or UPR induction (Figure 6 AC). This reinforces the idea that Atg39/40 are receptors for nutritional stress-induced and not constitutive ER-phagy.

We also tested if another constitutive ER-phagy cargo, Snq2-yEGFP, can be cleared by ERAD-C. Accumulation of overexpressed Snq2-yEGFP was compared in wild type, *trs85*Δ, *doa10*Δ, and *trs85*Δ *doa10*Δ mutant cells under normal growth conditions (SD+N). Unlike the effect on GFP-Snc1-PEM accumulation, *doa10*Δ had no effect on Snq2-GFP in combination with *trs85*Δ (Figure S5). This suggests that the partial overlap between constitutive ER-phagy and ERAD-C is cargo dependent (see below ERAD-M).

#### ERAD-C cargo and constitutive ER-phagy

We also wished to determine whether the ER-phagy pathway can contribute to clearance of an ERAD-C cargo, Deg1-Vma12-GFP. To that end, we tested whether *atg11*Δ, *trs85*Δ and *atg9*Δ are defective in clearance of Deg1-Vma12-GFP alone and in combination with *doa10*Δ using immuno-blot analysis. As was previously shown, WT cells do not accumulate Deg1-Vma12-GFP, but *doa10*Δ mutant cells do [29]. Like WT cells, single deletions of *ATG11, TRS85* and *ATG9* do not accumulate Deg1-Vma12-GFP. Interestingly, combining *doa10*Δ with *atg11*Δ, *trs85*Δ or *atg9*Δ results in an increase in accumulation of Deg1-Vma12-GFP above the level seen in the *doa10*Δ mutant cells to different levels, 30%, 2.5-fold and 3.5-fold, respectively (Figure 7A). In contrast, *atg39*Δ *atg40*Δ double-deletion mutant cells do not accumulate Deg1-Vma12-GFP, and do not increase the *doa10*Δ mutant phenotype (Figure 7B). These results show that while the constitutive ER-phagy proteins, Atg11, Trs85 and Atg9, can help with clearance on an ERAD substrate when Doa10 is absent, the nutritional stress-induced ER-phagy Atg39 and Atg40 receptors are not involved.

**Figure 7.**
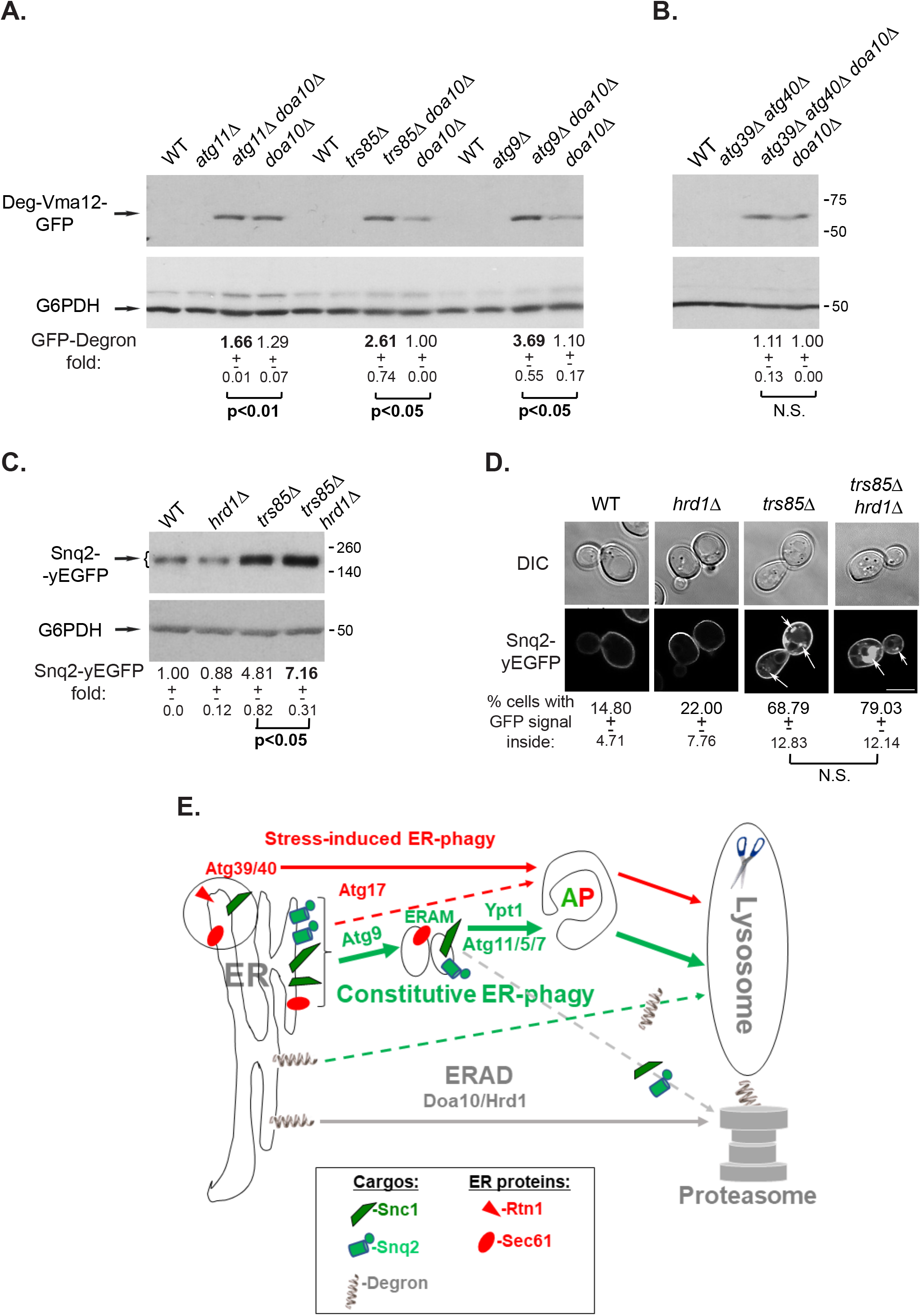
Partial overlap between constitutive ER-phagy and ERAD. **A-B.** Effect of constitutive ER-phagy mediators, but not of Atg39 and Atg40, on clearance of an ERAD-C cargo. **A.** Deletion of *ATG11, TRS85* and *ATG9* alone do not result in accumulation of an ERAD-C substrate, but they do increase the *doa10*Δ ERAD-C phenotype. WT and indicated mutant cells expressing Deg1-Vma12-GFP (integrated into the *TRP1* locus [29]) were grown in YPD. The level of Deg1-Vma12-GFP was determined by immuno-blot analysis using anti-GFP antibodies. Shown from top to bottom: strain, GFP blot, G6PDH blot (loading control), fold GFP-degron over the WT level, +/- STD, and p-value. Combining *atg11*Δ, *trs85*Δ or *atg9*Δ, with *doa1*Δ results in a significant increase of the GFP-Degron level. **B.**Deletion of *ATG39 ATG40* alone or in combination with *doa10*Δ does not result in accumulation of an ERAD-C substrate or an increase in the ERAD phenotype of *doa10*Δ. Experiments were done as in panel A. **C-D**. While deletion of *HRD1* alone does not result in Snq2-yEGFP accumulation, it does increase the defect of *trs85*Δ. WT and indicated mutant cells overexpressing Snq2-yEGFP were grown in normal growth medium (SD+N). The level of Snq2-yEGFP in cell lysates was determined by immuno-blot analysis using anti-GFP antibodies (**C**); intracellular accumulation of Snq2-yEGFP was determined by live-cell fluorescence microscopy (**D**). Results are presented as in Figure 2. A ~50% increase in the level of Snq2-yEGFP is observed when *hrd1*Δ is combined with *trs85*Δ. Results in this figure represent 3 independent experiments. **E**. Model showing the three ER quality control pathways discussed here: First, constitutive ER-phagy (green) shuttles overexpressed membrane proteins, e.g., Snc1 and Sec61, to the lysosome for degradation via autophagy. It requires, among other Atgs, Atg9, Ypt1, Atg11, Atg5 and Atg7. While Atg9 is required for sequestration of constitutive ER-phagy cargo into ER-associated membranes (ERAM), Ypt1 (and its GEF Trs85, not shown here), Atg11, Atg5 and Atg7 function in a later step of autophagosome (AP) assembly [7]. Second, Doa10- and Hrd1-mediated ERAD-C/M shuttle specific cargos, e.g., Deg1-Vma12-GFP, for degradation by the proteasome (gray). Both these processes operate during normal growth and depletion of each one does not affect the function of the other. However, when either constitutive ER-phagy or ERAD are compromised, the other process can clear some of the cargo (dashed lines). Third, nutritional stress-induced ER-phagy (red) shuttles ER fragments containing ER proteins, e.g., Rtn1, including constitutive ER-phagy cargo, e.g., Snc1 and Sec61, to the lysosome for degradation via APs. This process requires Atg17 and the Atg8-receptors Atg39 and Atg40. While these two receptors are not required for and do not affect neither constitutive ER-phagy nor ERAD, Atg17-dependent autophagy can clear constitutive ER-phagy cargo during normal growth in *atg11*Δ mutant cells (dashed red line).

#### Constitutive ER-phagy cargo and ERAD-M/L

Hrd1 is another ubiquitin ligase that recognizes misfolded regions in TMDs and lumenal domains of integral-membrane proteins, and facilitates their degradation via ERAD-M and ERAD-L, respectively [16]. We tested the effect of *hrd1*Δ on accumulation of two constitutive ER-phagy cargos, GFP-Snc1-PEM and Snq2-yEGFP, under normal growth conditions (SD+N) (Figures S6 and 7C-D, respectively). Like *doa10*Δ, *hrd1*Δ alone had no effect on accumulation of either cargo. Moreover, *hrd1*Δ did not affect the accumulation GFP-Snc1-PEM, when combined with *trs85*Δ, *doa10*Δ, or *trs85*Δ *doa10*Δ (Figure S6). In contrast, combining *hrd1*Δwith *trs85*Δ results in a 50% increase in the accumulation of Snq2-yEGFP (Figure 7 C-D). As for the partial overlap of constitutive ER-phagy and ERAD-C in the clearance of GFP-Snc1-PEM, the partial overlap of constitutive ER-phagy and ERAD-M in the clearance of Snq2-yEGFP was also apparent only when the protein level was quantified from an immuno-blot analysis and not by microscopy.

Together, our results show that ERAD-C and ERAD-M/L can partially help with clearance of cargo that accumulates in mutants defective in constitutive ER-phagy in a cargo-specific manner.

## Discussion

We show here that clearance of excess integral-membrane proteins during normal growth by macro-ER-phagy is constitutive and does not require induction of general autophagy. Moreover, while the two AIM receptors, Atg39 and Atg40, are required for nutritional stress-induced ER-phagy [15], including clearance of excess integral-membrane proteins (shown here), they neither play a role in constitutive ER-phagy nor serve as its cargos. Interestingly, while Atg17 is not required for clearing excess integral-membrane proteins during normal growth, it can fulfill this role when Atg11 is deleted. This shows that the general autophagy machinery can partially compensate for the Atg11-dependent constitutive ER-phagy. Lastly, we determine that while disruption of ER-phagy or ERAD does not perturb the other process, disruption of each one exacerbates defects of the other. This shows that these two ER-quality control processes that function during normal growth can partially compensate for each other (Figure 7E).

### Constitutive versus nutritional stress-induced ER-phagy

While ER-phagy of overexpressed integral-membrane proteins occurs under normal growth conditions, it was possible that excess of integral-membrane proteins stimulates stress-induced autophagy. Five findings support the idea that this is not the case and that this process is constitutive. First, increase in Atg8 level is a hallmark of nutritional stress-induced autophagy [23]. Although clearance of excess integral-membrane proteins during normal growth requires Atg8, and proteins needed for its lipidation, Atg5 and Atg7, Atg8 level is not increased. Second, measuring Pho8Δ60 activity is the accepted quantitative nutritional stress-induced autophagy assay [24]. During normal growth, neither overexpression of an integral-membrane protein nor its accumulation in mutants defective in its clearance, result in an induction of Pho8Δ60 activity. Third, Atg11, which is known to be required for another constitutive autophagy pathway, CVT [6], is required delivering excess integral-membrane proteins to the vacuole during normal growth, whereas Atg17, which is required for nutritional stress-induced autophagy [12], is not required for this process. Fourth, processing of the CVT cargo, Ape1, is affected by starvation both in wild-type cells (from partial to complete processing) and in *atg11*Δ mutant cells (from complete block to partial processing). Overexpression of an integral-membrane protein during normal growth does not affect Ape1 processing neither in wild type nor in *atg11*Δ mutant cells. Finally, the two AIM receptors required for nutritional stress-induced ER-phagy, Atg39 and Atg40, are not required for ER-phagy during normal growth.

In summary, constitutive ER-phagy uses the machinery of general macro-autophagy. The three differences between constitutive and nutritional stress-induced ER-phagy are: First, constitutive ER-phagy does not need the specific receptors of stress-induced selective-ER-phagy (Atg39, Atg40). Second, it does not need to be induced by nutritional or ER stress. Third, constitutive and stress-induced ER-phagy use mostly Atg11 and Atg17, respectively. Notably, in the absence of Atg11, constitutive ER cargo can be shuttled to the lysosome in an Atg17-dependent manner. Thus, we conclude that ER-phagy during normal growth is a constitutive process and delivery of endogenously-expressed cargo can be further induced by overexpression of an integral-membrane protein.

### What is the cargo for constitutive ER-phagy?

During normal growth, constitutive ER-phagy cargos are delivered to the vacuole in wild-type cells and accumulate together outside the vacuole in the cytoplasm of mutant cells defective in constitutive ER-phagy. Previously, we have shown that overexpressed integral-membrane proteins that are normally transported to the PM, GFP-Snc1-PEM, DsRed-Snc1-PEM and Snq2-yEGFP, serve as cargos for constitutive ER-phagy. In addition, two ER resident proteins, Sec61 and Hmg1, are delivered to the vacuole for degradation through this pathway even without overexpression of membrane proteins. In contrast, two proteins that mediate formation of ER-to-Golgi vesicles, the membrane protein Sec12 and the coat protein Sec13, do not serve as cargos for constitutive ER-phagy even when GFP-Snc1-PEM is overexpressed [7]. Here, we add three ER resident proteins that do not serve as constitutive ER-phagy cargos, the two nutritional stress-induced ER-phagy receptors, Atg39 and Atg40, and Rtn1, delivery of which to the vacuole under nutritional stress depends on Atg40. Interestingly, in *ypt1-1* mutant cells, regions of Rtn1-labeled and Snc1-labeled ER seem to be adjacent, but separate (Figure 4C). These results suggest that separate ER domains contain proteins that during normal growth can serve as cargos for constitutive ER-phagy and domains that do not contain them. Distinct ER domains are thought to perform the multiple roles of the ER [31].

### What happens to cargo of constitutive ER-phagy when the cells are stressed?

Under nutritional stress, Atg17 and the two receptors, Atg39/40, play a major role in ER-phagy [12, 15]. In wild-type cells under nutritional stress, constitutive ER-phagy cargo can be shuttled to the vacuole in an Atg17- and Atg39/40-dependent way like other ER proteins. The same is true for constitutive ER-phagy cargo that accumulates in mutant cells defective in this process, e.g., *atg11*Δ, which can be at least partially cleared in an Atg17, Atg39, and Atg40-dependent way. Thus, under nutritional stress, all regions of the ER, including those that can serve as constitutive ER-phagy cargo, can be shuttled through stress-induced selective ER-phagy for degradation in the vacuole.

### Partial overlap between constitutive ER-phagy and ERAD

Another ER-QC pathway that functions during normal growth is ERAD, which shuttles misfolded proteins to the proteasome for degradation. Topologically, the cargo for ERAD-C is similar to that of GFP-Snc1-PEM, a constitutive ER-phagy, namely a transmembrane protein with a large cytoplasmic domain [16]. The question was if the two processes can recognize the other’s cargo. Depletion of components required for ERAD-C or constitutive ER-phagy alone do not result in defects in processing the other’s cargo. This suggests that each system is potent enough to clear its own substrate. However, depletion of both ERAD-C and constitutive ER-phagy components results in more severe defects in processing of both cargos suggesting partial overlap of the two processes.

ERAD-C cargo that accumulates in a mutant defective in this process during normal growth can be cleared by constitutive ER-phagy. Specifically, combining *doa10*Δ with either *atg11*Δ, *trs85*Δ, or *atg9*Δ, results in increased accumulation of the ERAD-C cargo Deg1-Vma12-GFP. Moreover, the ER-phagy receptors Atg39 and Atg40 are not required for this clearance. Thus, constitutive ER-phagy can deliver ERAD-C substrates to the vacuole.

Clearance of GFP-Snc1-PEM, a constitutive ER-phagy cargo, by ERAD-C depends on the nature of its accumulation. Specifically, an increase in the unprocessed cargo of constitutive ER-phagy GFP-Snc1-PEM was seen when *doa10*Δ was combined with *atg11*Δ, *trs85*Δ or *atg5*Δ, but not with *atg9*Δ. We have previously showed that Atg9 functions before Atg11 and Trs85 in constitutive ER-phagy. Whereas constitutive ER-phagy cargo accumulates in the ER of *atg9*Δ mutant cells, in *atg11*Δ and *trs85*Δ it accumulates in aberrant ER structures we termed ER-to-autophagy membranes (ERAM) [7]. Therefore, Doa10 seems to recognize excess of integral-membrane protein that accumulates in ERAM but not in the ER.

Interestingly, ERAD-C does not recognize a second constitutive ER-phagy cargo, Snq2-yEGFP. Because in addition to a cytoplasmic domain, this protein has multiple TMDs, we asked whether it can be recognized by ERAD-M. The E3 Ub ligase that mediated ERAD-M/L is Hrd1 [16]. While *hrd1*Δ does not cause Snq2-yEGFP accumulation, it does increase its accumulation in *trs85*Δ mutant cells by 50%. Conversely, *hrd1*Δ does not exacerbate the accumulation phenotype of GFP-Snc1-PEM of *trs85*Δ or *trs85*Δ *doa10*Δ. Thus, Snc1 and Snq2 that accumulate in mutants defective in constitutive ER-phagy, can be partially cleared by ERAD-C and ERAD-M/L, respectively. Therefore, the partial suppression of constitutive ER-phagy defects by ERAD-C or ERAD-M/L is cargo dependent.

Together, these results show that in addition to misfolded proteins, ERAD can recognize excess proteins, and constitutive ER-phagy can recognize misfolded proteins in addition to excess proteins and send them for degradation by the proteasome and the vacuole, respectively (Figure 7E).

### Does UPR play a role in clearance of constitutive ER-phagy cargo

Accumulation of excess or misfolded proteins in the ER results in ER stress and induction of the unfolded protein response (UPR) [19]. Whereas UPR is constitutively induced in mutants defective in intracellular traffic, it is not induced in mutants defective only in autophagy [26]. We have previously shown that in most mutants defective in autophagy, overexpression of GFP-Snc1-PEM results in UPR induction. However, UPR induction is not required for constitutive ER-phagy to occur [7]. Here, we show that clearance of constitutive ER-phagy cargo by ERAD-C also does not require UPR induction. Thus, while UPR is induced when excess integral-membrane protein accumulates in the ER, it does not seem to affect its clearance by either constitutive ER-phagy or ERAD-C.

### Is there an overlap between constitutive- and micro-ER-phagy?

Micro-ER-phagy can be induced by ER stress [22] or overexpression of certain ER resident proteins [32]. As for cargo, at least one cargo, Sec61, can be shuttled to the vacuole via both constitutive- and micro-ER-phagy [7, 32]. This is expected because micro-ER-phagy delivers big regions of the ER [22, 32]. However, the two pathways do not overlap. First, Atg proteins are not required for micro-ER-phagy [22, 32], and we showed that Ypt1 is also not required for it [7]. Second, because the destination of both macro- and micro-ER-phagy is the vacuole, if micro-ER phagy could shuttle GFP-Snc1-PEM to the vacuole (in addition to constitutive ER-phagy), the accumulation of this cargo is expected to be higher when vacuolar proteases are depleted than when just macro-ER-phagy is defective. However, *pep4*Δ *prb1*Δ mutant cells accumulate similar levels of GFP-Snc1-PEM to that in mutants defective in constitutive ER-phagy (e.g., *ypt1-1*). Third, unlike ERAD, micro-ER-phagy does not shuttle to the vacuole constitutive ER-phagy cargo that accumulates in mutants defective in this process. This is supported by the observation that depletion of vacuolar proteases (Pep4 and Prb1) in such mutants (e.g., *ypt1-1*) does not result in additional accumulation of the cargo [7].

### Comparison of ER proteostasis in yeast and mammalian cells

As in yeast cells, misfolded proteins are cleared from the ER in mammalian cells by ERAD and ER-to-lysosome associated degradation (ERLAD) [33]. Three different mammalian ERLAD pathways were described. First, micro-ER-phagy was shown to remove misfolded procollagen in mouse osteoblasts [34]. Second, ERAD-resistant polymers are delivered from the ER to lysosomes via vesicles and not APs [35]. Third, misfolded (mutant) procollagen is cleared via a selective macro-ER-phagy that requires the FAM134B receptor [36], a suggested mammalian counterpart of the yeast Atg40 (based on domain architecture resemblance [15]). The constitutive ER-phagy pathway described here is different from all three mammalian ERLAD pathways. As discussed above, we ruled out micro-ER-phagy and AP-independent delivery to the lysosome. While the mammalian selective macro-ER-phagy of misfolded collagen is the closest to the constitutive-ER-phagy described here, there is no evidence that proteins need to be misfolded for serving as cargo for the latter pathway, and the selective ER receptors Atg39 and Atg40 are not required for constitutive ER-phagy in yeast. Finally, the known cargos the three mammalian ERLAD pathways are lumenal, while the cargos for constitutive ER-phagy in yeast is integral-membrane proteins. Therefore, the question whether excess integral-membrane proteins are cleared from the ER by constitutive macro-ER-phagy in mammalian cells is still open.

In summary, we characterize here clearance of excess membrane protein during normal growth as constitutive ER-phagy. Importantly, this process is not dependent on a known stress, nutritional or misfolded protein, or response to this stress. We also show that while constitutive ER-phagy is different from nutritional stress-induced ER-phagy, which clears ER indiscriminately, it can partially overlap with it, e.g., in *atg11*Δ, Atg17-dependent autophagy can clear constitutive ER-phagy cargo. It also partially overlaps with another ERQC process ERAD, which clears misfolded proteins. These findings raise the following new questions for future research. First, what is the mechanism by which constitutive ER-phagy cargo is recognized during normal growth? Second, if cargo sorting for constitutive ER-phagy depends on ER domain organization, what is the basis for organization of the different ER domains: ER-to-Golgi transport (Sec12), constitutive ER-phagy cargos (Sec61, Hmg1), perinuclear ER (Atg39), and ER underlying the PM (Atg40, Rtn1)? Third, what is the basis for the partial overlap between constitutive ER-phagy and ERAD? Finally, what about constitutive ER-phagy in mammalian cells? Constitutive autophagy was implicated in cell homeostasis [37], differentiation [38], and aging [39]. Moreover, accumulation of overexpressed membrane proteins is relevant to human disease, notably in inflammation and cancer [40-43]. Furthermore, protein overexpression can lead to aggregation [44], which in turn contribute to neurodegenerative disorders [45], diabetes [46] and cancer [47]. However, even though autophagy in general and nutritional stress-induced ER-phagy specifically are conserved between yeast and human cells [48], not much is known about mechanisms of constitutive autophagy of any kind in human cells.

## Materials and Methods

### Strains, plasmids and reagents

Yeast (*Saccharomyces cerevisiae*) strains used in this paper are summarized in Table S1. Plasmids used in this study are summarized in Table S2. Gene disruptions and tagging were performed by a standard PCR-based method ([49]. All chemical reagents were purchased from Fisher Scientific (NJ, USA), except for the following: Nitrogen bases were purchased from US Biological (Swampscott, MA); ProtoGel for Western blots from National Diagnostics (Atlanta, GA); Bacto peptone and Bacto agar from BD Difco (Franklin Lakes, NJ); salmon testes DNA, amino acids, 1-naphthyl phosphate, 2-nitrophenyl β-D-galactopyranoside and protease inhibitors from Sigma Aldrich (St. Louis, MO); glass beads from BioSpec Products (Bartlesville, OK); restriction enzymes and buffers from New England Biolabs (Ipswich, MA).

Antibodies used in this study included mouse monoclonal anti-GFP (Roche Diagnostics), rat anti-HA (Roche Diagnostics), mouse anti-HA (Covance), rabbit anti-G6PDH (Sigma), rabbit anti-Ape1 (a kind gift from Dr. Ohsumi), rabbit anti-Atg8 (a kind gift from Dr. Ohsumi), goat anti-rat HRP (Abcam), goat anti-rabbit HRP and goat anti-mouse HRP (GE Healthcare).

### Plasmid construction

Rtn1 and Nop1 were tagged with GFP on the chromosome. Genomic DNA fragments containing native promoter, ORF, and GFP tag with ADH1 terminator were amplified and cloned in pRS313 using EcoRV/BmgBI and XbaI for Rtn1 and EcoRV/BmgBI and XbaI/NheI for Nop1.

### Yeast culture conditions

Medium preparation and yeast culture growth for nitrogen-starvation shift experiments were done as described [50]. For Atg39/40 experiments cells grown to mid-log phase at 26°C were treated with 200 ng/ml rapamycin for 14 to 18 hours.

### Protein level analyses

To determine levels of GFP-tagged proteins in yeast lysates cell pellets were resuspended in water then vortexed with the addition of 0.26N sodium hydroxide and 0.13M beta-mercaptoethanol. Proteins were precipitated by the addition of 5% trichloroacetic acid then suspended in SDS sample buffer and boiled for 10 min. Preparation of protein lysates for Ape1 processing analysis and determination of Atg8 level was done as described [10]. Urea sample buffer was used for protein level analysis of Atg39 as described except the cells were disrupted by vortexing with 0.5mm glass beads [15]. The following protease inhibitors were added to the lysis buffers: 10 μg/ml leupeptin, 10 μg/ml aprotinin, 10 μg/ml chymostatin, 5 μg/ml pepstatin A, 1 mM benzamidine, 5 μg/ml antipain, 0.1 mM PMSF. ImageJ was used for quantification of protein bands; when comparing levels between lanes, the levels were normalized using a loading control (G6PDH). For assessing degradation of a GFP-tagged protein under stress, e.g., -N or +rapamycin, the percent of the full-size protein degraded to GFP is determined. Under normal growth, free GFP does not accumulate (the protein is completely degraded, including the GFP moiety). GFP-Snc1-PEM protein runs as a doublet (shown by a bracket) due to phosphorylation [51]; both bands were used for the quantification. A degradation block during normal growth is determined by an increase in the level of the fullsize protein in autophagy-defective mutants compared to wild-type cells. In wild type cells, decrease in level of full-length protein is due to degradation in the vacuole because it is reversed in mutants defective in vacuolar proteases, e.g., *pep4*Δ *prb1*Δ. In autophagy-defective mutants, depletion of vacuolar proteases does not affect the level of the protein because the block is before the protein reached the vacuole [7].

### ALP activity and UPR β-gal activity assays

Alkaline phosphatase activity assay of Pho8Δ60 was done as previously described [52]. The UPR β-gal assay uses a plasmid expressing LacZ behind four UPR elements (UPRE, 22 bp from *KAR2* promoter) [53, 54] and was done as previously described [7].

### Microscopy

Live cell microscopy was done as follows: wild-type and mutant cells carrying constructs for expression of GFP or mCherry-tagged protein(s) were grown to mid-log phase in appropriate selective media. Fluorescent microscopy was performed using a deconvolution Axioscope microscope (Carl Zeiss) with FITC filter set or confocal LSM700 microscope (Carl Zeiss) controlled by Zen software using 488 nm (GFP) and 555 (mCherry) laser lines. Colocalization analysis was carried out using the ImageJ software: Specifically, images were preprocessed to minimize noise and background, the color threshold values were manually adjusted, and co-localization rates were determined by calculating the area of co-localization compared to the total fluorescence of the region of interest. Accumulation of GFP-Snc1-PEM in mutant cells was quantified and expressed in two different ways: 1) during normal growth, most of the GFP in wild-type cells is on the PM, and “percent cells with GFP signal inside” was quantified. 2) Under nutritional stress, in wild type cells, GFP can reach the vacuole through ER-phagy, therefore, to distinguish accumulation in wild type and mutant cells “percent cells with GFP signal inside (not in vacuole)” (vacuolar membrane is labeled with FM4-64) was determined.

### Statistical analyses

Unpaired TTEST analysis was applied for determining statistical significance for experiments with three repeats or for a pair of samples. The significance of p values are shown as: n.s (not significant) for p>o.05; or p<0.05 is considered significant.

## Acknowledgments

We thank Dr. Yoshinori Ohsumi for the kind gift of anti-Atg8 antibodies, and Dr. Mark Hochstrasser and Dr. Jason Berk for the Deg1-Vma12-GFP strains.

## Supplementary Information

Tables S1-S2

Figures S1-S6

## Supplemental Figures

**Figure S1.**
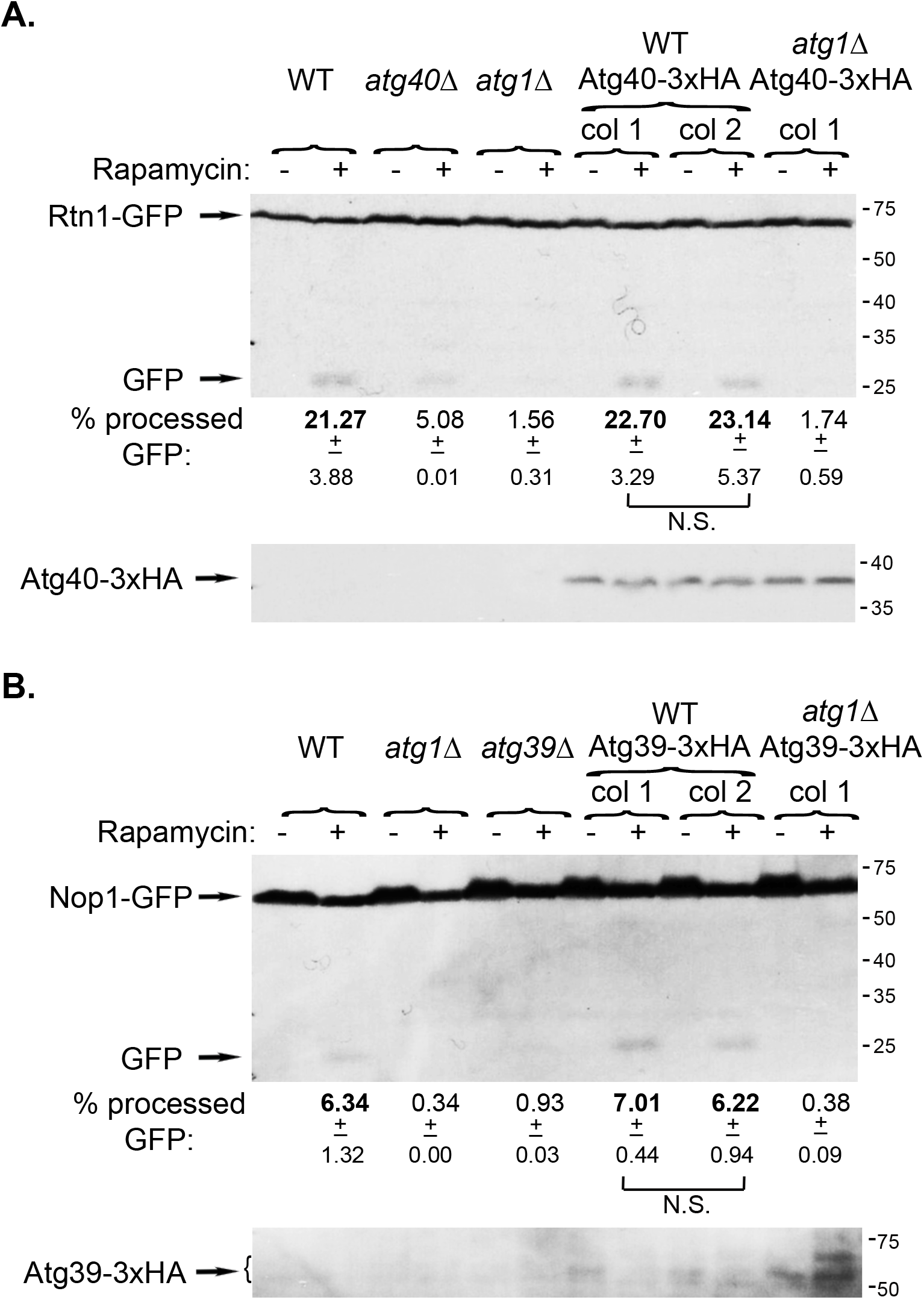
HA-tagged Atg40 and Atg39 are functional. **A.** Atg40-3xHA can shuttle its Rtn1-GFP ER cargo for degradation under rapamycin-induced stress. Wild type and *atg1*Δ mutant cells expressing endogenously tagged Atg40-3xHA were transformed with a CEN plasmid expressing Rtn1-GFP under its own promoter. Cells were grown in normal medium (YPD) or under stress (+Rapamycin) and processing of Rtn1-GFP to GFP was determined by immunoblot using anti-GFP antibodies. From left to right: WT, *atg40*Δ, *atg1*Δ, WT expressing Atg40-3xHA (two colonies), and *atg1*Δ expressing Atg40-3xHA. Shown from top to bottom: strain, growth conditions (- or + rapamycin), GFP blot, quantification of the % processed GFP, +/- STD, p-value, and HA blot (to confirm expression of Atg40-3xHA). >20% of the Rtn1-GFP is processed to GFP under stress in WT, but not *atg40*Δ or *atg1*Δ mutant cells. Importantly, cells expressing Atg40-3xHA as the only copy process Rtn1-GFP as well as WT cells (but not in *atg1*Δ cells). **B.** Atg39-3xHA can shuttle its Nop1-GFP ER cargo for degradation under stress. The same experiment described in panel A, was done with cells expressing endogenously tagged Atg39-3xHA and transformed with a CEN plasmid expressing Nop1-GFP under its own promoter. ~6-7% of the Nop1-GFP is processed to GFP under stress in WT, but not *atg39*Δ or *atg1*Δ mutant cells. Importantly, cells expressing Atg39-HA (bands within the bracket [15]) as the only copy, process Nop1-GFP as well as WT cells (but not in *atg1*Δ cells). Note: the level of Atg39-HA is >25-fold lower when cells are grown in YPD (as was done in this experiment that did not require selection of a plasmid) than in SD [55]. Results in this figure represent 3 independent experiments.

**Figure S2.**
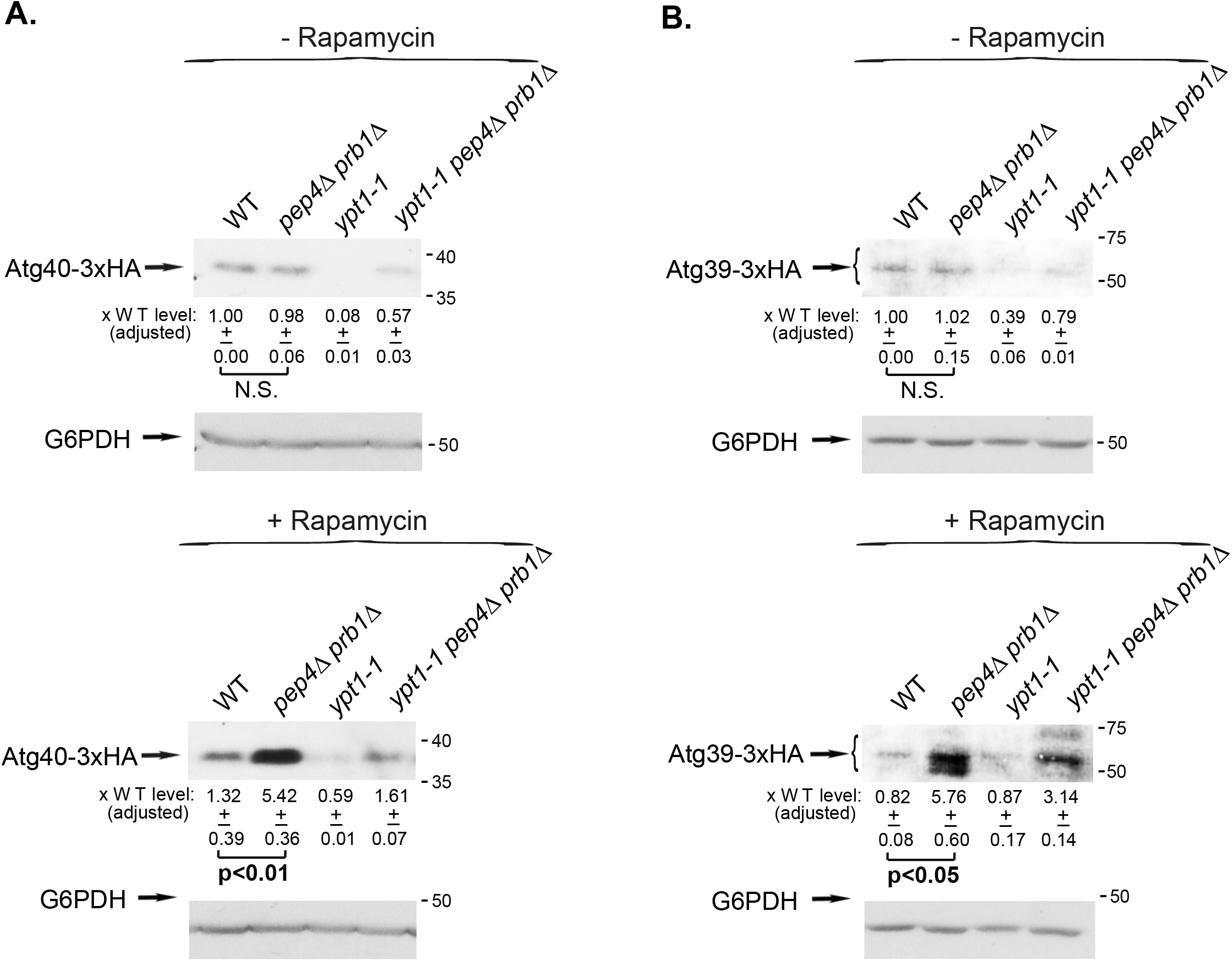
HA-tagged Atg40 and Atg39 levels in *ypt1-1* and protease deficient mutant cells. Endogenous Atg40 **(A)** and Atg39 **(B)** were tagged with 3xHA at their C-termini in four strains, WT, *pep4*Δ *prb1*Δ, *ypt1-1*, and *ypt1-1 pep4*Δ *prb1*Δ. Cells were grown in normal medium (SD-Rapamycin, top) or under stress (SD+Rapamycin, bottom). The level of Atg40-3xHA and Atg39-3xHA in cell lysates was determined using anti-HA antibodies and immunoblot analysis (G6PDH was used as a loading control). Shown from top to bottom: growth conditions, strain, HA blot, quantification of the HA-tagged protein band: fold over wt, +/- STD, p value, and G6PDH blot. Under stress (+Rapamycin), the levels of both Atg40-3xHA and Atg39-3xHA (bands within the bracket [15]) increases by ~5-folds in cell defective in vacuolar proteolysis (*pep4*Δ *prb1*Δ), showing that they are delivered for degradation to the vacuole. The levels of Atg40 and Atg39 are lower in *ypt1-1* mutant cells during normal growth and they increase during nutritional stress in proteolysis defective cells. Results from in this figure represent 3 independent experiments.

**Figure S3.**
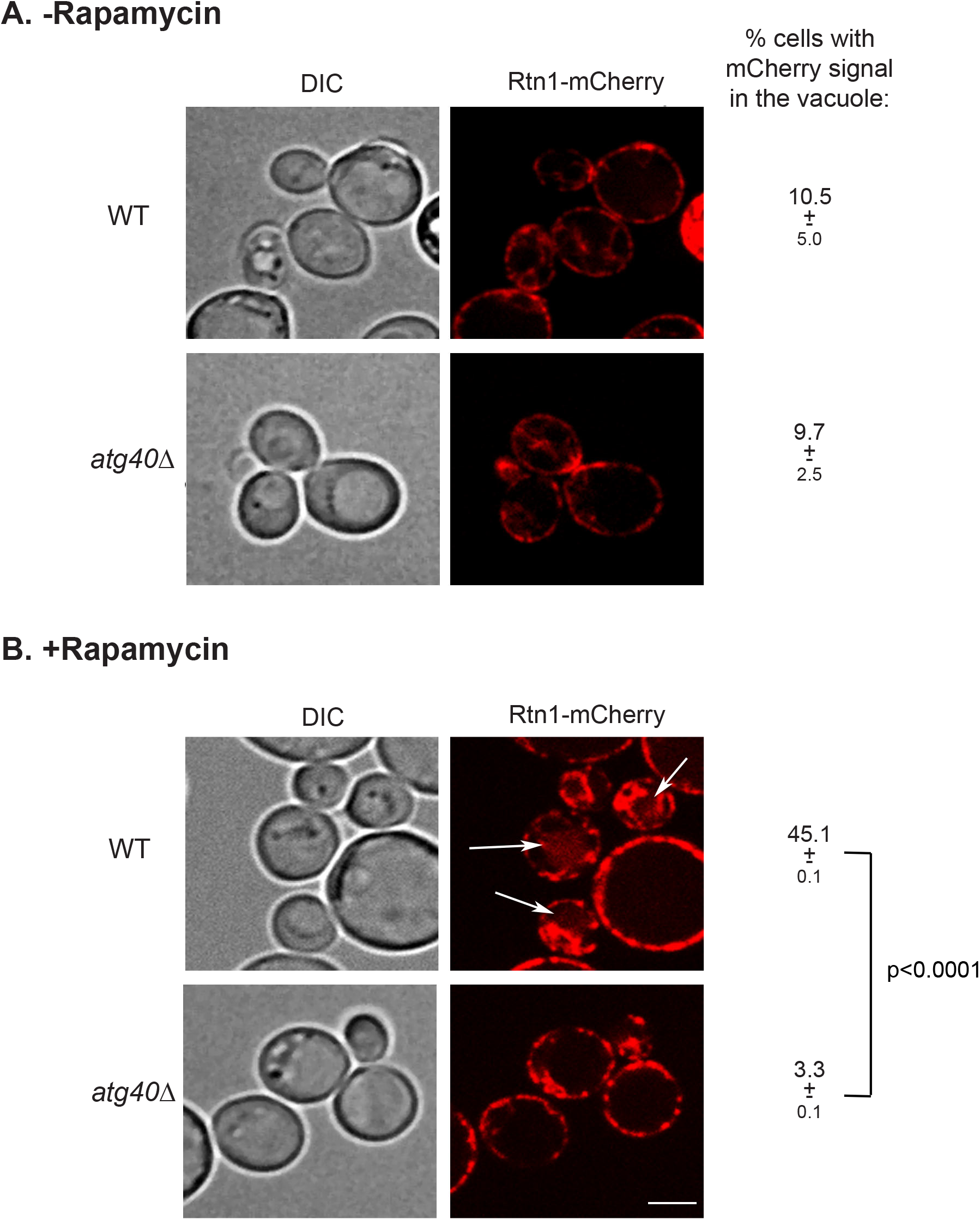
Rtn1-mCherry is an Atg40 cargo. Endogenous Rtn1 was tagged with mCherry at its C-terminus (as in Figure 4C) in wild type and *atg40*Δ mutant cells. Cells were grown to mid log (A) and treated with rapamycin for 16 hours (B), were visualized by live-cell fluorescence microscopy. Shown from left to right: DIC, mCherry, % cells with Rtn1-mCherry in the vacuole; +/-, STD, and p value. Rtn1-mCherry localizes to the ER of both wild type and *atg40*Δ mutant cells during normal growth. Under stress (+rapamycin), it is delivered to the vacuole in 45% of wild type, but not *atg40*Δ mutant, cells. >150 cells were visualized for each data point; arrows point to Rtn1-mCherry in the vacuole; size bar, 1μ. Results in this figure represent 4 independent experiments.

**Figure S4.**
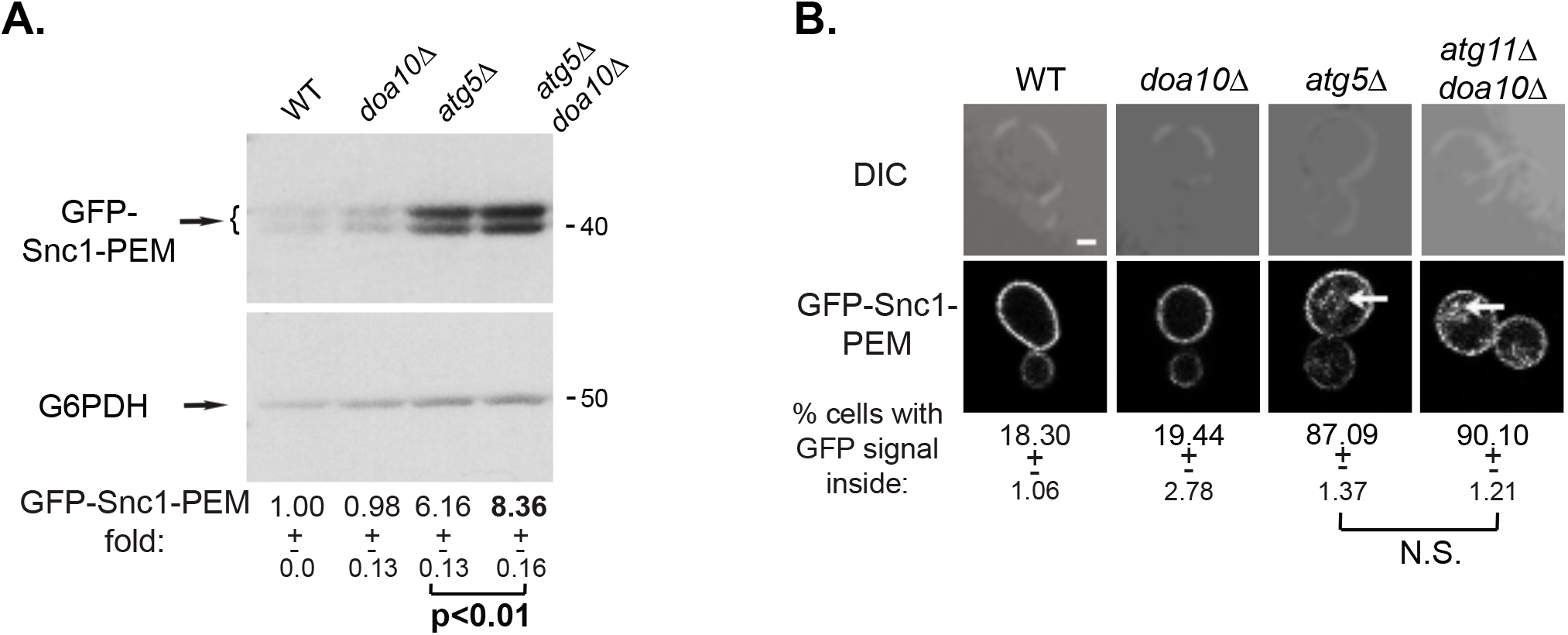
Deletion of *DOA10* increases GFP-Snc1-PEM accumulation in *atg5*Δ. WT and indicated mutant cells overexpressing GFP-Snc1-PEM were grown in normal growth medium (SD+N). The level of GFP-Snc1-PEM in cell lysates was determined by immuno-blot analysis using anti-GFP antibodies (**A**); intracellular accumulation of GFP-Snc1-PEM was determined by live-cell fluorescence microscopy (**B**). Results are presented as in Figure 2. A 35% increase in the level of GFP-Snc1-PEM is observed when *doa10*Δ is combined with *atg5Δ.*Results in this figure represent 3 independent experiments.

**Figure S5.**
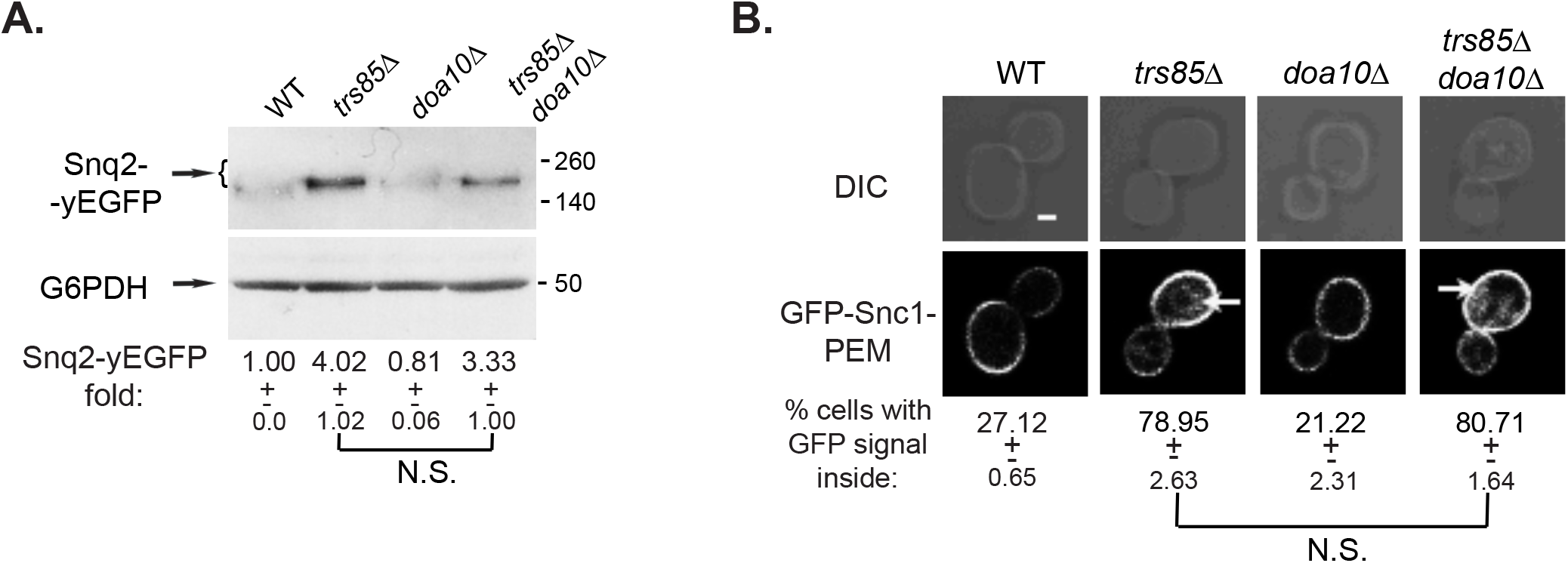
ERAD-C does not affect clearance of the constitutive ER-phagy cargo Snq2-yEGFP. WT and indicated mutant cells overexpressing Snq2-yEGFP were grown in normal growth medium (SD+N). The level of Snq2-yEGFP in cell lysates was determined by immunoblot analysis using anti-GFP antibodies (**A**); intracellular accumulation of Snq2-yEGFP was determined by live-cell fluorescence microscopy (**B**). Results are presented as in Figure 2. No increase in the level of Snq2-GFP is observed in *doa10*Δ or when it is combined with *trs85*Δ. Results in this figure represent 3 independent experiments.

**Figure S6.**
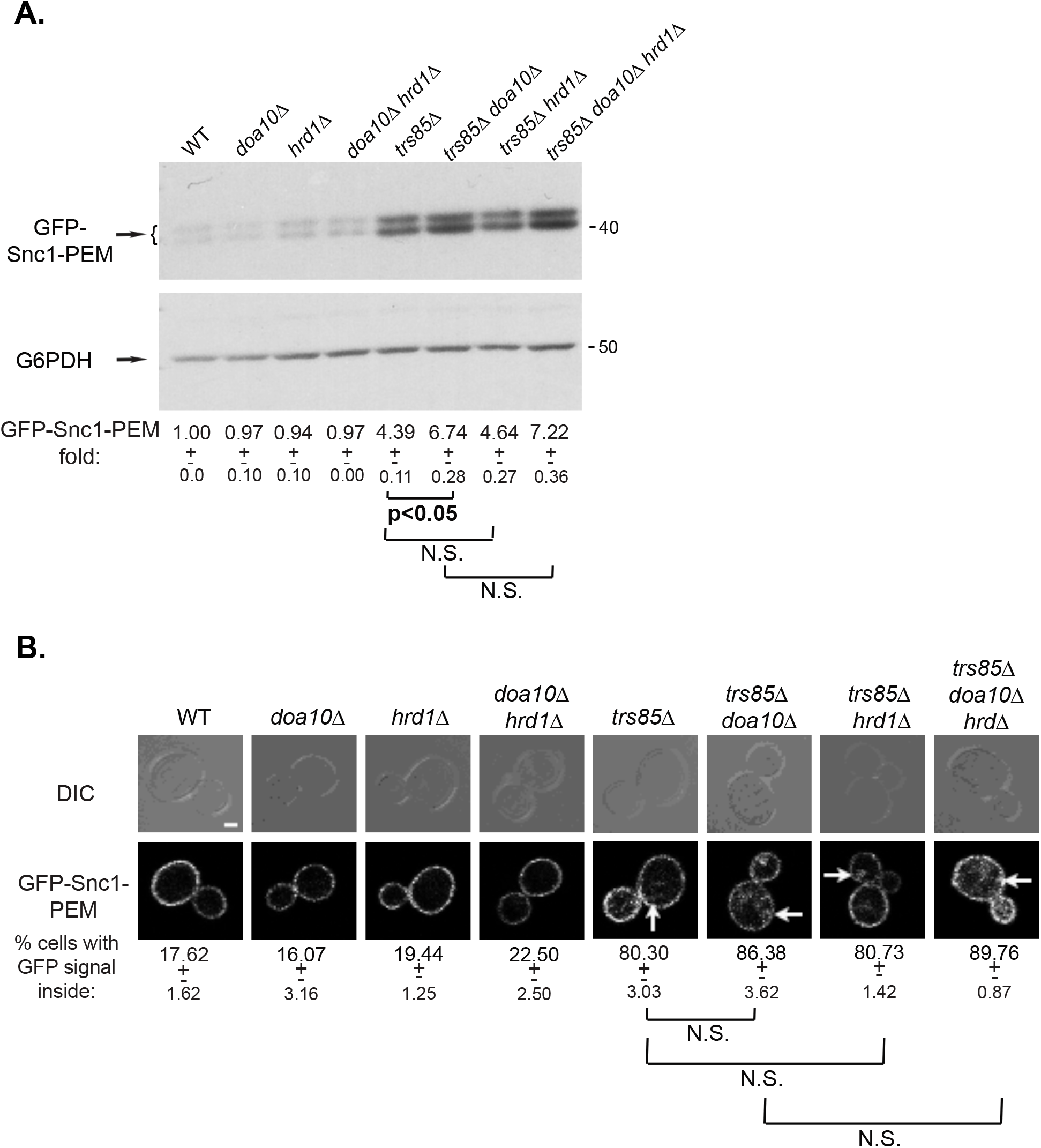
ERAD-M/L does not affect clearance of the constitutive ER-phagy cargo GFP-Snc1-PEM. WT and indicated mutant cells overexpressing GFP-Snc1-PEM were grown in normal growth medium (SD+N). The level of GFP-Snc1-PEM in cell lysates was determined by immuno-blot analysis using anti-GFP antibodies (A); intracellular accumulation of GFP-Snc1-PEM was determined by live-cell fluorescence microscopy (B). Results are presented as in Figure 2, Whereas *doa10*Δ results in 50% increase of GFP-Snc1-PEM when combined with *trs85*Δ, no increase in the level of GFP-Snc1-PEM is observed in *hrd1*Δ, or when it is combined with *trs85*Δ, *doa10* or *trs85*Δ *doa10*. Results in this figure represent 2 independent experiments.

---

**Table S1.**
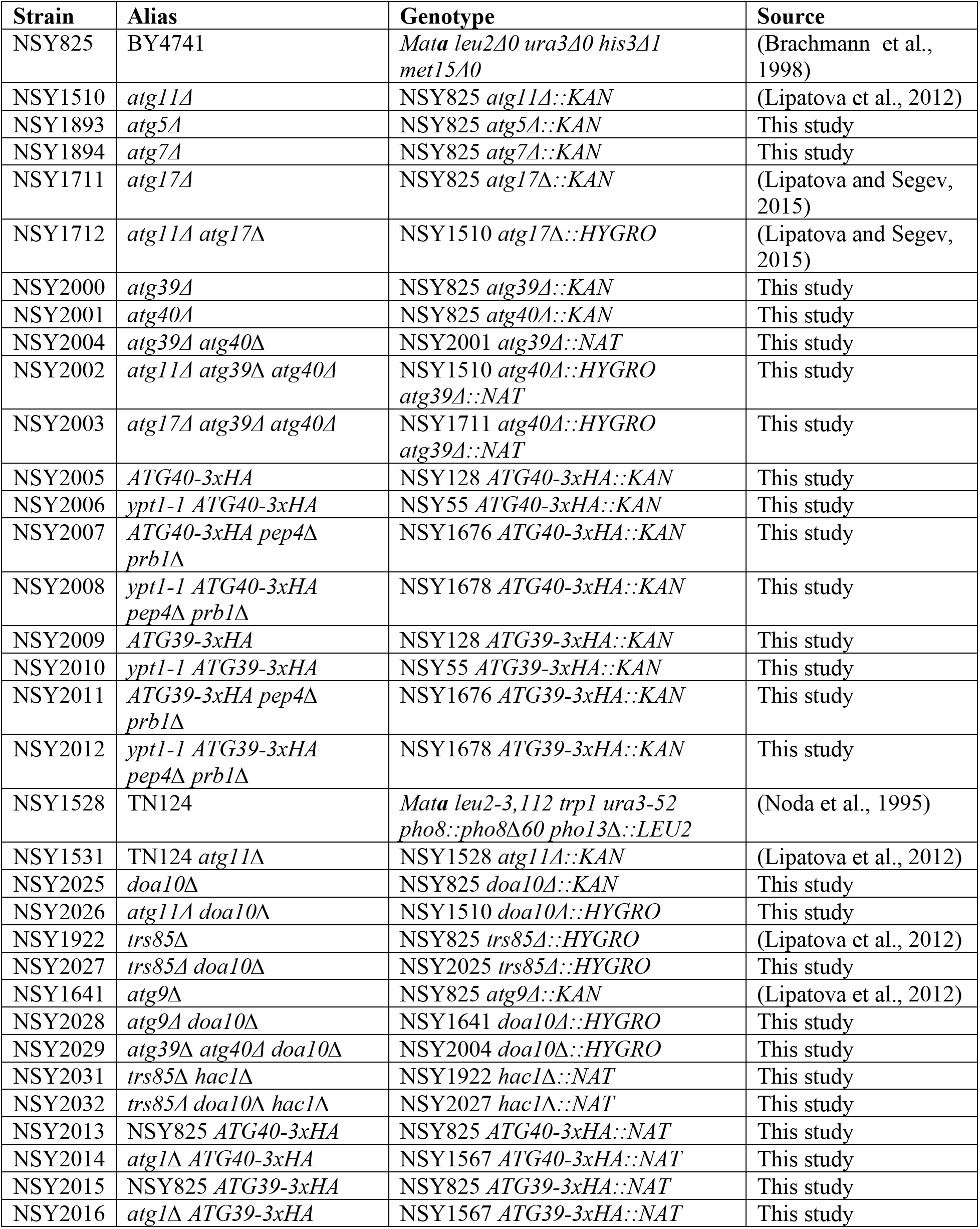
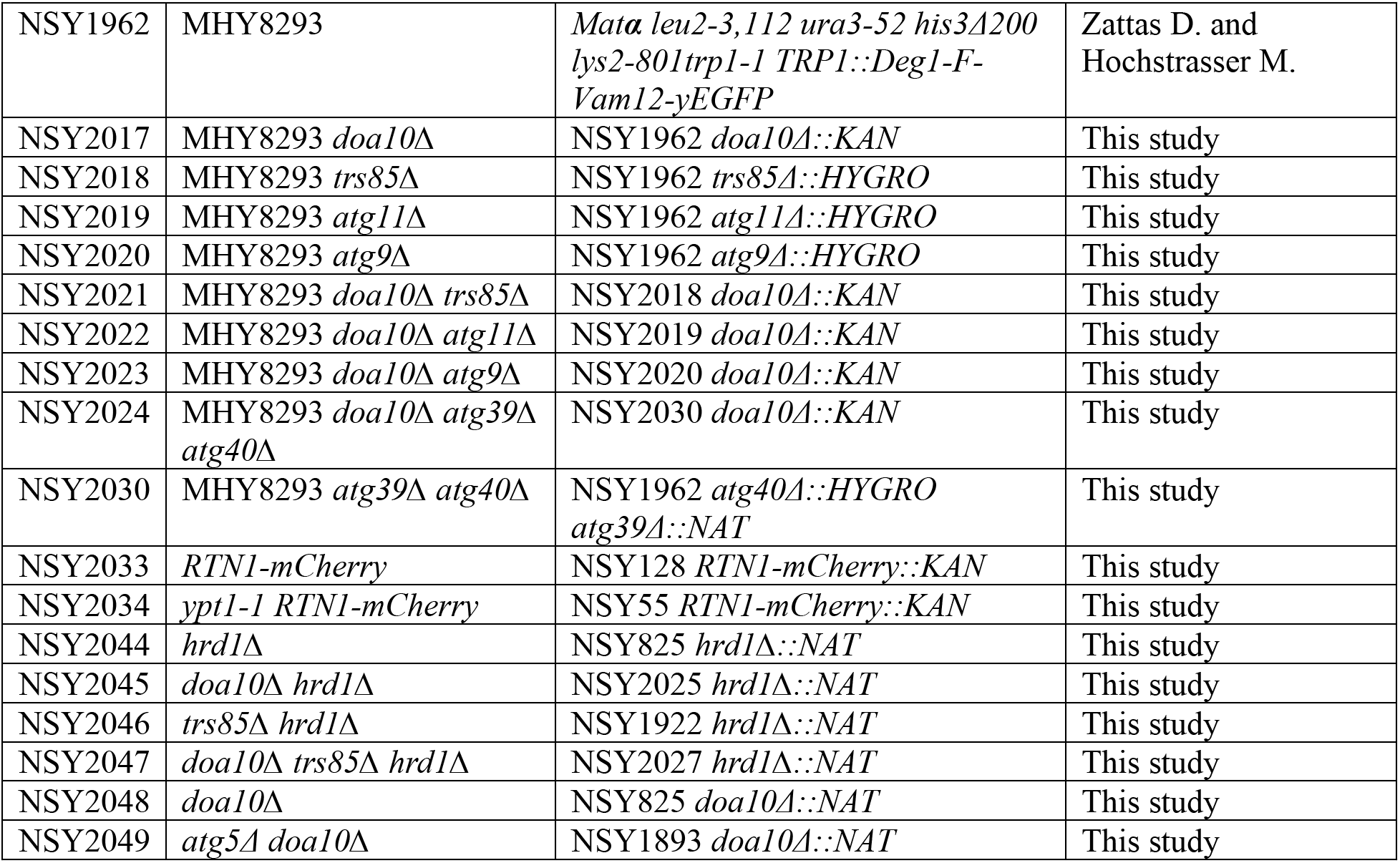
Yeast strains used in this study.

**Table S2.**
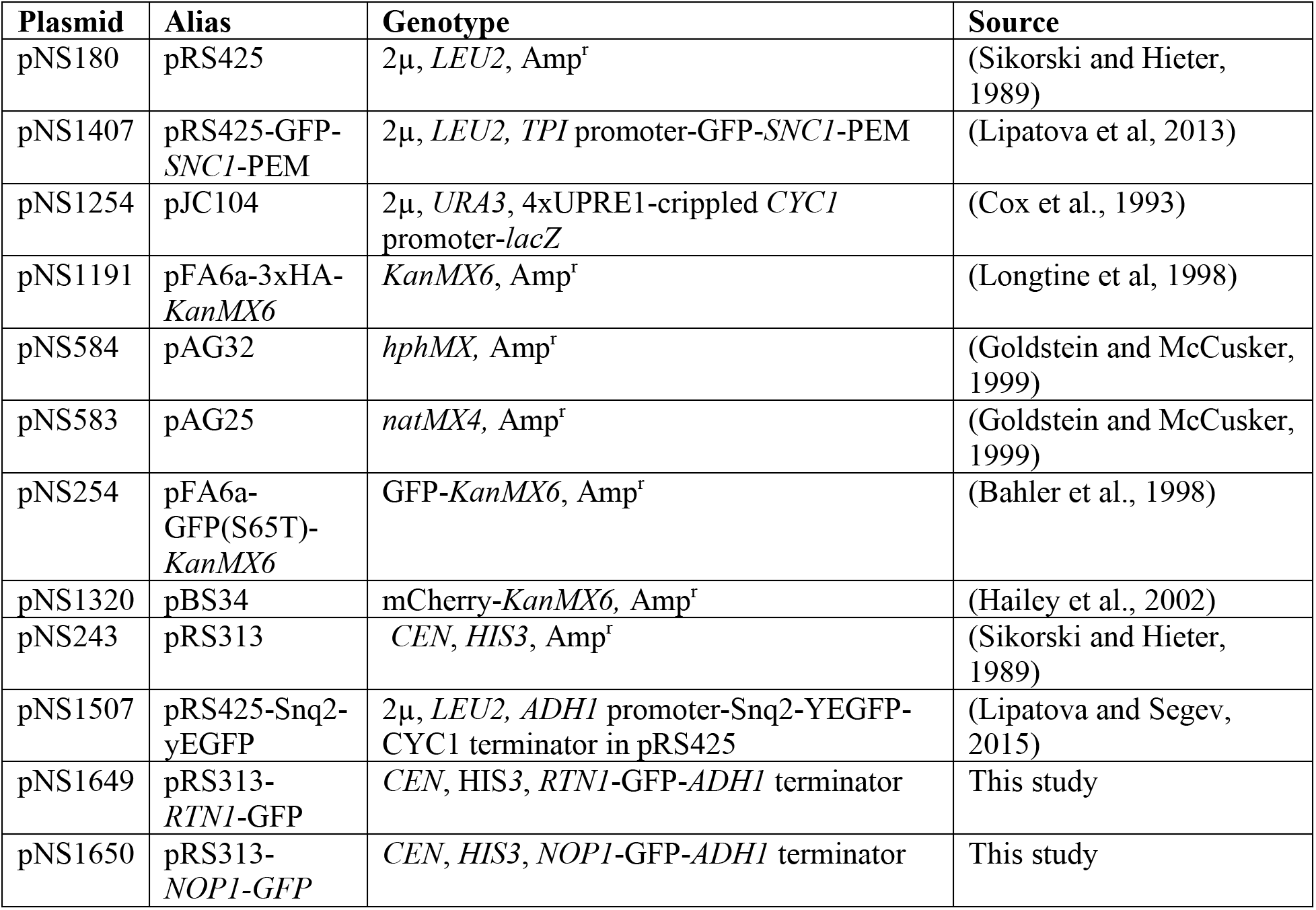
Plasmids used in this study.

